# A novel expression system for imaging single-molecule fluorescence in Haloferax volcanii WR806 enables visualization of Cas1 involvement in UV-light associated DNA repair

**DOI:** 10.1101/2025.02.24.639823

**Authors:** Paula R Schrage, Uliana Afonina, Julia Wörtz, Anita Marchfelder, Koen JA Martens, James P Sáenz, Ulrike Endesfelder

## Abstract

Fluorescence microscopy has become an indispensable tool in biological research, offering powerful approaches to study protein dynamics and molecular biochemistry *in vivo*. Among archaea, *Haloferax volcanii* has emerged as a particularly well-suited model organism for imaging studies, with a growing toolkit of established fluorescent markers, plasmids, and promoter systems. Recent advances in single-molecule imaging techniques have created new opportunities through WR806, a carotenoid-free strain providing reduced autofluorescence background. However, existing plasmid-based expression systems in WR806 show critical limitations in protein expression control and challenges with protein aggregation.

To address these limitations, we developed pUE001, a novel expression system specifically designed for WR806. This system achieves precise expression control by decoupling selection and induction through strategic implementation of the *trpA* selection marker. Through comprehensive characterization, we demonstrate that pUE001 provides superior control over protein expression compared to the previously established pTA962 system. It enables linear, titratable expression of diverse proteins — from the highly regulated CRISPR-Cas component Cas1 to the abundant structural protein FtsZ1 — while preventing protein aggregation that could compromise native cellular functions. Additionally, we performed a comprehensive analysis of WR806 to show that carotenoid depletion does not affect native cellular physiology. Finally, to demonstrate the system’s utility, we investigated the role of Cas1 in UV-induced DNA repair using single-particle tracking photoactivated localization microscopy (sptPALM). Our findings reveal significant, dose-dependent changes in Cas1 mobility following UV-light induced damage, providing evidence for its involvement in DNA repair processes and offering new insights into the expanding roles of CRISPR-Cas systems beyond adaptive immunity.

## INTRODUCTION

Fluorescence microscopy has emerged as a powerful tool for cell biological research, enabling the study of protein dynamics, subcellular localization, and cellular biochemistry *in vivo*. Recent advances in tools for both diffraction-limited and super-resolution microscopy have established imaging as an essential method in archaeal research, providing direct visual insights into cellular processes (van Wolferen *et al*. 2022). For *Haloferax volcanii*, a model organism for halophilic archaea, there is a quickly growing toolkit available, e.g. fluorescent proteins (FPs) (Turkowyd *et al*. 2020; Ithurbide *et al*. 2024), plasmid systems (Allers *et al*. 2004; Ithurbide *et al*. 2024) and regulated promoters (Large *et al*. 2007; Rados *et al*. 2023), which has facilitated investigations into cell shape dynamics (Duggin *et al*. 2015; Walsh *et al*. 2019), DNA replication (Delpech *et al*. 2018) and cell division (Liao *et al*. 2021; Nußbaum *et al*. 2021, 2024).

As live-cell fluorescence microscopy becomes increasingly central to cell biology research, maintaining native cell biology while fluorescently labelling molecules of interest has emerged as a critical challenge. Two key issues have been reported for *H. volcanii*: First, genetic modifications, such as the deletion of essential genes for auxotrophic selection, can trigger unexpected metabolic effects, as demonstrated in the pleomorphic archaeon (Patro *et al*. 2023). Second, expressing genes from non-endogenous loci such as plasmid systems based on pHV2 (seemingly advantageous due to *H. volcanii’s* polyploid nature) can result in non-native transcription and protein production, potentially disrupting native biology through various mechanisms, including morphological variance (Patro *et al*. 2023), impaired cell division (Liao *et al*. 2021), altered protein dynamics, and the formation of protein aggregates such as inclusion bodies (Fahnert, Lilie and Neubauer 2004). Consequently, minimizing these artefacts requires thoughtful selection of strains, growth conditions, and expression systems.

Recently, our group established single-molecule localization microscopy (SMLM) and single-particle tracking (SPT) methods for *H. volcanii*. In this proof-of-concept work, we super-resolved FtsZ1 ring structures and tracked RNA polymerase dynamics in living *H. volcanii* cells (Turkowyd *et al*. 2020). A significant challenge arose from *H. volcanii*’s natural production of colorful carotenoids like bacterioruberin, generating high cellular autofluorescence and interfering with the detection of single fluorescent molecules in SMLM imaging. To overcome this limitation, we generated a carotenoid-free “albino” strain WR806 by disrupting the biosynthesis pathway through deletion of the *crtI* gene. Additionally, we established codon-optimized versions of two photoswitchable fluorescent proteins using the tryptophan-inducible pTA962 plasmid expression system: PAmCherry1Hfx and Dendra2Hfx (Turkowyd *et al*. 2020).

Despite its advantages for single-molecule imaging, two critical issues emerged. First, although bacterioruberin is relatively scarce in *H. volcanii*, constituting less than 5% of cell membrane content (Kellermann *et al*. 2016), carotenoids in general have been shown to play a distinct role for different bacterial membranes, such as protection against oxidative stress (Kim *et al*. 2019; Rizk *et al*. 2021), alteration of membrane fluidity by increased production at lowered temperatures (Seel *et al*. 2020) or contributing to light scavenging in phototrophs (Polívka and Frank 2010). Even though WR806 exhibits wild-type-like growth (Turkowyd *et al*. 2020), the impact of carotenoid depletion on its membrane biophysics and consequent physiological effects in *H. volcanii* remained to be comprehensively investigated. Second, the plasmid system pTA962 presented significant challenges in controlling protein expression when used in WR806. We e.g. found that protein aggregation and variable expression levels of proteins of interest (POIs) can compromise studies of native cell biology (unpublished preliminary data). The use of plasmid-based systems for expressing modified proteins (e.g. fluorescently-labeled, mutated, or truncated), is typically faster to implement experimentally when compared to genomic integration. While WR806’s multiple auxotrophic markers enable flexible selection strategies, they also create unique constraints for plasmid expression systems. Specifically, the strain carries deletions in *trpA* (tryptophan synthesis, tryptophan synthetase alpha subunit), *hdrB* (thymidine synthesis), *leuB* (leucine synthesis), and *pyrE2* (uracil synthesis) genes (Ortenberg, Rozenblatt-Rosen and Mevarech 2000; Bitan-Banin, Ortenberg and Mevarech 2003; Allers *et al*. 2004). While these deletions enable experiments based on multiple plasmids, they necessitate – if not complemented by plasmids carrying the corresponding synthesis genes - external supplementation of tryptophan, thymidine, and uracil (leucine supplementation can be typically neglected as present in standard growth media). Consequently, effective plasmid systems for WR806 must satisfy two key criteria: compatibility with this auxotrophic background and controlled protein expression through inducible promoters to minimize disruption of native cell biology.

Currently, *H. volcanii* has only two available inducible promoter systems: p.tna, the native promotor of the tryptophanase gene *tnaA* (HVO_0009) (Large *et al*. 2007) and p.xyl-750, the native promoter of the xylose degradation operon *xacEA* (HVO_B0027-28) (Rados *et al*. 2023). Due to high leaky expression observed with p.xyl-750 (Rados *et al*. 2023), we focused on the rather tightly controlled tryptophan-inducible p.tna system (Large *et al*. 2007). However, our previously used pTA962 plasmid (Turkowyd *et al*. 2020), presents a significant limitation when used in WR806: while pTA962 complements *hdrB* and *pyrE2* selection markers in WR806, it lacks *trpA* complementation needed for tryptophan synthesis, rendering WR806 carrying pTA962 dependent on external tryptophan supply in the medium. The direct tryptophan uptake from the medium creates an inherent problem as the plasmid’s tryptophan-inducible p.tna promoter (originally designed for strains with genomic *trpA* for internal synthesis, (Allers *et al*. 2010)) is induced even at the minimal tryptophan concentrations required for growth. This unintended coupling of selection and induction leads to constitutive POI expression from pTA962. Two strategies can address this issue: 1) engineering a *crtI*-gene-deficient mutant of a *trpA*-encoding strain like H26 or H98 (Allers *et al*. 2004), or 2) incorporating *trpA* directly onto the plasmid used for POI expression. We preferred the latter approach, since it maintains minimal use of selection markers.

The native promoter of the *trpCBA* operon is not tightly regulated (Large *et al*. 2007). At the same time, expression levels of POI under a p.tna promoter are tightly regulated in strains carrying genomic *trpCBA* (Large *et al*. 2007). We thus hypothesized that providing a plasmid-based copy of *trpA* under the constitutive p.fdx promoter would result in a similar, post-transcriptional, control of tryptophan synthesis as for the native *trpCBA* operon. To test this, we developed pUE001, a novel expression plasmid that combines the backbone from pTA231 (encoding the *trpA* selection marker under the control of p.fdx, (Allers *et al*. 2004)), with the p.tna promoter region from pTA962 (Allers *et al*. 2010). As we were able to show in this work, this system effectively decouples induction from selection, providing improved control over protein expression levels. Furthermore, by design, pUE001 utilizes only the *trpA* selection marker, flexibly preserving WR806’s remaining auxotrophic markers (*hdrB*, *leuB*, and *pyrE2*) for additional genetic manipulations.

To summarize, this work makes two key contributions: First, we present a detailed comparative analysis of WR806 demonstrating that carotenoid depletion does not affect native cellular processes, particularly membrane biophysics. Second, we establish that our novel plasmid pUE001, combined with the carotenoid-free strain WR806, creates an ideal tool for low-background fluorescence imaging in *H. volcanii*. Finally, we validate this by successfully tracking the single-molecule dynamics of Cas1, a highly regulated, low-abundant protein of the type I-B Clustered Regularly Interspaced Short Palindromic Repeats and CRISPR-associated genes (CRISPR-Cas) system in *H. volcanii*.

## MATERIAL AND METHODS

### Strains, media, and cultivation conditions

The strains used in this study are listed in Supplementary Table 1.

*E. coli* strains were grown over night at 37 °C in LB liquid culture shaking at 180 rpm or on LB Agar plates with 1.5 % (w/v) agar-agar. Agar plates and liquid cultures were supplied with a final concentration of 100 µg/mL of carbenecillin for selection of plasmids.

All *H. volcanii* strains were grown at 42°C in liquid HV-Cab medium (Duggin *et al*. 2015; De Silva *et al*. 2021) or HV-YPC medium. HV-Cab agar plates with 1.5 % (w/v) agar-agar were used to streak out strains from cryostocks or for plating after transformation. The additives uracil, thymidine and tryptophan were added in final concentrations of 0.45 mM, 0.16 mM, and 0.25 mM, respectively. WR806 and H119 were supplied with additional tryptophan, thymidine and uracil (Supplementary Table 1). WR806 carrying pTA962 plasmid constructs were supplied with tryptophan, while pUE001 carrying strains were supplied with tryptophan and uracil (Supplementary Table 1). Induction of the p.tna promoter system was accomplished by supplying strains with tryptophan in concentrations between 0.25 and 5 mM.

### Determination of growth rates

Growth rates of H119 and WR806 were measured using a Tecan Infinite® M PLEX plate reader. Single colonies were selected from agar plates, and cultures were inoculated in either HV-Cab or HV-YPC medium and grown to stationary phase. Cultures were then diluted to an OD_600_ of 0.1 and transferred into a 48-well plate. Growth curves were monitored over 72 hours at 42 °C, with brief shaking performed every 20 minutes before measurements were taken.

### Construction of plasmids

Vector constructs were created by Gibson assembly using the NEBuilder® HiFi DNA Assembly Cloning Kit (NEB, Frankfurt am Main, Germany) according to the manufacturer’s protocol. All oligonucleotides to amplify backbones and inserts are listed in Supplementary Table 2, all plasmids are listed in Supplementary Table 3. Backbones and inserts were amplified by PCR with Q5® High Fidelity DNA Polymerase (NEB, Frankfurt am Main, Germany). To construct the plasmids pUE001-FtsZ1:Dendra2Hfx (Addgene # 234671) and pUE001-Cas1:Dendra2Hfx (Addgene # 234670), the plasmid pTA231-p.Syn-Dendra2Hfx was used as template for backbone amplification. To amplify the promoter and terminator region, including the coding sequence for the fusion proteins FtsZ1:Dendra2Hfx and Cas1:Dendra2Hfx, the plasmids pTA962-FtsZ1:Dendra2Hfx and pTA962-Cas1:Dendra2Hfx were used as templates. Consecutively, pUE001-Cas1:Dendra2Hfx was used to construct pUE001-Dendra2Hfx (Supplementary Figure S1, Addgene # 234669). To reduce the number of original template plasmids prior to transformation, all products were digested using DpnI (NEB, Frankfurt am Main, Germany).

### Strain creation

Vector constructs were transformed into competent *E. coli* DH5α cells via heat-shock to ensure efficient plasmid replication. Plasmid sequences were verified using Sanger and Oxford Nanopore sequencing (Eurofins genomics, Cologne, Germany). The plasmids were then passaged through *E. coli dam-/dcm-* to obtain non-methylated plasmid DNA, which was subsequently transformed into *H. volcanii* WR806 using PEG600 (Cline et al., 1989). Transformed cells were streaked onto HV-Cab agar plates containing the respective selection additives (Supplementary Table 3) Positive transformants were checked by colony PCR (Supplementary Table 2).

### Lipid extraction

Lipid extracts used for absorbance spectra, FTIR spectra and monolayer assays (Figure 1 and Supplementary Figure S2) were obtained from H119 and WR806 grown in from single colonies in HV-Cab medium to stationary phase. Lipid extraction was performed a standard protocol (Bligh and Dyer 1959) with modifications. Briefly, Cells were pelleted at 6000 rpm for 15 min and separated into 25 mg pellets. Pellets were washed with PBS and cell lysis was achieved by sonication for 20 min. Subsequently, 300 µL chloroform:methanol (1:1, v/v), 100 µL 0.1 M HCl, and 100 µL 0.5 M NaOH were added. Samples were shaken vigorously for 5 min, followed by 40 min centrifugation at 9000 rpm. Again, 200 µL of 0.1 M and 0.5 M NaOH were added, respectively, and samples were shaken vigorously for 15 min. Lastly, after centrifugation for 3 min at 13000 rpm, the organic, lower phase was removed from the sample and solvents were evaporated under nitrogen flow. Lipids were stored at -20 °C until further analysis.

**Figure 1.**
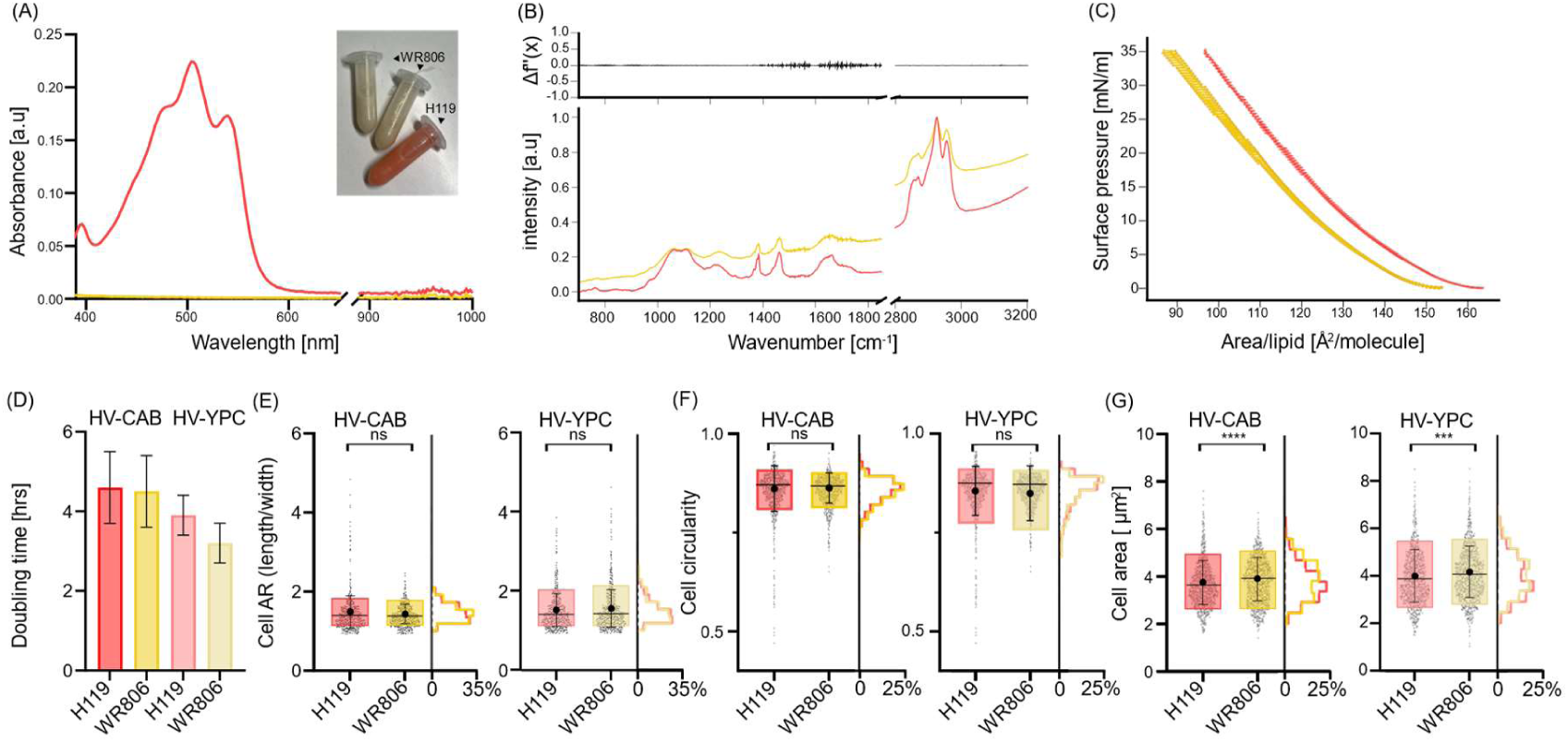
Comparative characterization of *H. volcanii* strains H119 and WR806. Strain H119 (DS70-wildtype (ΔpHV2), *ΔpyrE2*, *ΔtrpA*, *ΔleuB*; (Allers *et al*. 2004)) was compared to the carotenoid-deficient “albino” strain WR806 (DS70-wildtype (ΔpHV2), *ΔpyrE2*, *ΔtrpA*, *ΔleuB*, *ΔhdrB*, *ΔcrtI*; (Turkowyd *et al*. 2020)), with *ΔcrtI* deletion being responsible to disrupt carotenoid biosynthesis. Data from H119 and WR806 are shown in red and yellow, respectively. (A) UV-VIS absorbance spectra (technical duplicates) with inset showing Eppendorf tubes containing total lipid extracts; (B) FTIR spectra, normalized by intensity, of lipid extracts (technical duplicates; lower panel) and their second-order derivative differences (upper panel); (C) Area isotherms measuring surface pressure versus molecular area for lipid monolayers at 42°C (two biological replicates, each with five technical replicates); (D) Growth rates expressed as doubling times at 42°C in HV-Cab and HV-YPC media (means ± std. dev.; two biological replicates, each with three technical replicates); (E-G) Quantitative morphological analysis of cell populations by microscopy, showing distributions of (E) aspect ratio, (F) circularity, and (G) cell area per single cell (n > 950 cells per strain). Box plots display 10-90% quartiles with mean (big circle), median (line) and standard deviation; histograms show frequency distributions. Significance by Mann-Whitney-test. ns: not significant; ****: P < 0.0001; ***: P = 0.0002.

### Phospholipid quantification

The concentration of phospholipids was quantified as described in (Katewa and Katyare 2003). Briefly, phosphorus ICP standard solution (Merck, Germany) at concentrations of 0, 5, 10, 50, and 100 nmol was used to determine the standard curve. Samples and standards were mixed with 500 µL 70% HClO_4_, mixed thoroughly and incubated at 200 °C for 2 hrs. After 30 min cooling, 500 µL 10 % (w/v) ascorbic acid (stock solution in dH2O) and 500 µl 2.5 % (w/v) (NH_4_)_6_Mo_7_O_24_ · 4 H_2_O (stock solution in dH2O) was added to each sample, mixing in between steps. After 30 min incubation at 37 °C, absorbance of each sample was measured at 820 nm wavelength in a plate reader. Concentrations of lipid extracts were calculated based on the standard curve, which was plotted as linear fit of the A_820_ function of moles of inorganic phosphate.

### Lipid monolayer assay

Lipid monolayer assays were measured at 37 °C and 42 °C using a Langmuir Blodgett trough Microtrough G1 (Kibron, Finland) and using the lipid extracts from H119 and WR806 cultures. Chloroform solutions of lipid extracts were directly prepared at the concentrations determined by the phosphate assay. Monolayers were formed by injecting 13 µL and 30 µL of lipid solution onto an aqueous subphase (buffer of 3.3 mM sodium citrate, 3.3 mM sodium phosphate, 3.3 mM glycine and 0.15 M NaCl at pH 7.5) maintained at 37°C and 42 °C, respectively, using a built-in temperature-controlled circulating water bath. Isotherms were recorded with a 70 cm^2^ Teflon Langmuir trough, which was equipped with a motorized compression barrier and a pressure sensor (Kibron DeltaPi, Finland). The area per lipid was calculated by dividing the total lipid area by the number of lipid molecules as determined by a phosphate quantification assay (Katewa and Katyare 2003). For each mixture, the area per lipid was estimated from the averages of isotherms obtained from two monolayers, with three technical replicates for 37 °C and five replicates for 42 °C. The areas corresponding to the closest pressure values to the defined pressure points (5, 10, 15, 20, 25, 30, and 35 mN/m) were selected for the averaged isotherms. The compressibility modulus (k) (Supplementary Figure S2C, D) represents the membrane’s resistance to compression and was also calculated using the Langmuir monolayer experimental data. Compressability is defined as: k = -A * (∂π/∂A)T where A is the molecular area, π is the surface pressure, and the subscript T indicates the derivative is taken at constant temperature (Przykaza *et al*. 2019). This dimensionless parameter quantifies how the molecular area changes in response to applied surface pressure.

### FTIR-Spectroscopy

For FTIR spectroscopy measurements of lipid extracts from H119 and WR806, the samples were embedded into KBr optical windows. First, 50 µL sample was mixed with 250 mg of finely ground, dry KBr powder. The dried sample was pressed under 8T pressure for 5 min. FTIR spectra were recorded on an INVENIO Tensor spectrometer (Bruker, USA), with a KBr window used as the background. All spectra were processed and analyzed using the OPUS software (Bruker, USA), with normalization and peak searching performed by custom Python code.

### Plate reader experiments

Plate reader experiments on a Tecan Infinite® M PLEX were performed at 42°C for time-course measurements of OD_600_ and fluorescence intensity and in dependency of tryptophan concentration used for induction of the p.tna promoter. Single colonies were picked from HV-Cab agar plates and inoculated in selective HV-Cab medium until stationary phase was reached. Prior to the plate reader run, cultures were diluted to OD_600_ 0.1 with final tryptophan concentrations between 0 and 5 mM. All samples were measured in biological triplicates with technical duplicates each in Greiner µClear® CELLSTAR® black 96 well plates. Measurements were taken every 20 minutes after a brief orbital shaking step of 4 mm for 30 s. OD_600_ measurements were blanked using the medium values from the respective runs. Fluorescence was measured at an excitation wavelength of 470 nm and emission window of 510 ± 9 nm matching green Dendra2Hfx fluorescence. The gain was determined for each POI individually. Fluorescence measurements were blanked to the average fluorescence of WR806 WT as a negative control over time. For obtained the normalized fluorescence intensity curves of Figure 2, measured fluorescence intensities were then divided by the respective OD measurement, to normalize to the cell count.

**Figure 2.**
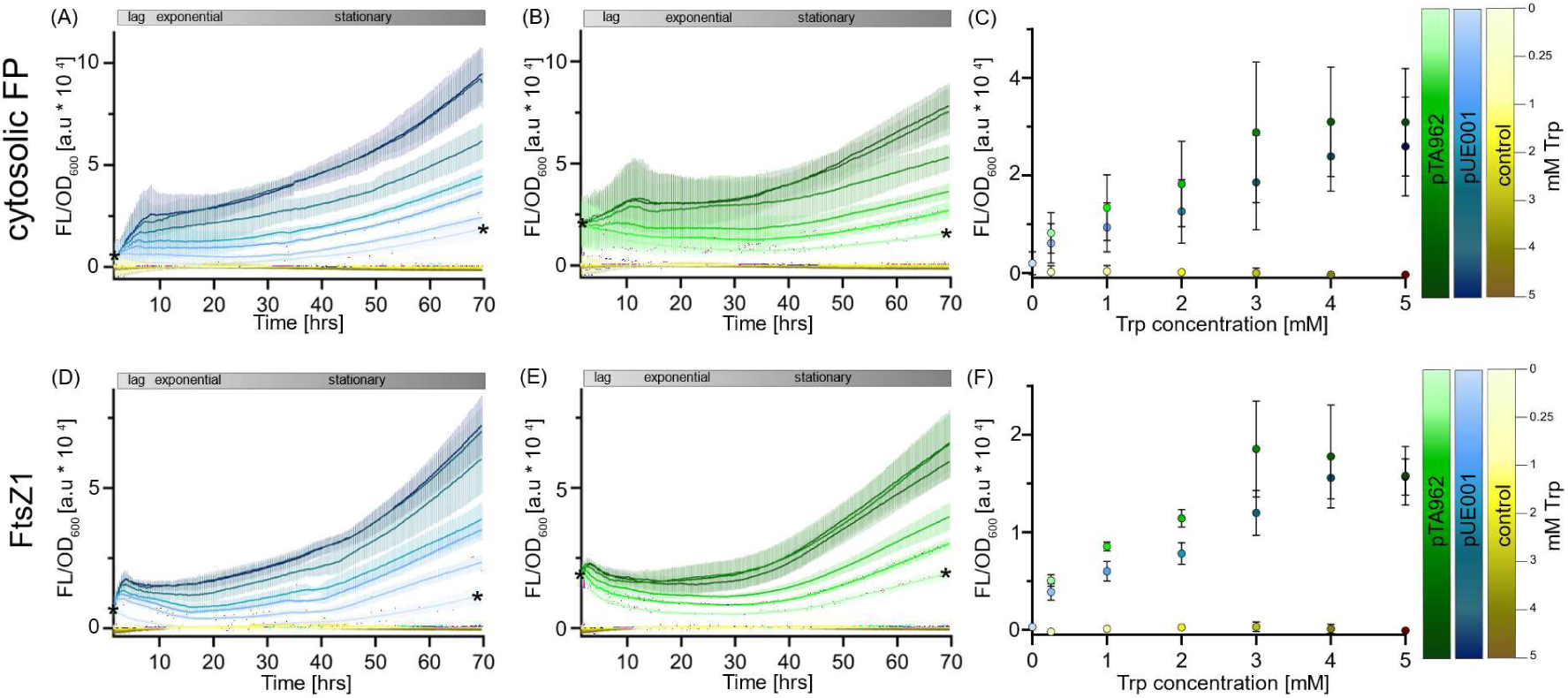
Fluorescence characterization of Dendra2Hfx expression in *H. volcanii* strain WR806. (A-F) Time-course fluorescence measurements of tryptophan-induced protein expression normalized to cell density OD600 (ex 470 nm; em 501-519 nm, optimized for Dendra2Hfx green fluorescence). Measurements were performed at 42°C in biological triplicates, each with technical duplicates, data is visualized as mean with standard deviation. (A, B) Cytosolic Dendra2Hfx expression from pTA962 (green) and pUE001 (blue) plasmids (plasmid details in Supplementary Table S2; Supplementary Figure S1). (D, E) FtsZ1:Dendra2Hfx fusion protein expression from the same plasmid systems. Cultures were induced with tryptophan concentrations of 0, 0.25, 1, 2, 3, 4, and 5 mM. Plasmid-free WR806 (yellow) served as negative control. Initial fluorescence reflects protein accumulation from autoinduction of three-day stationary phase pre-cultures. Asterisks at 0 and 70h indicate the matching fluorescence levels of initial values and stationary phase cultures. Growth phases (lag, exponential, and stationary) were determined from raw data of OD600 measurements (Supplementary Figure S3). (C, F) Steady-state fluorescence levels during exponential growth phase, with cell growth and protein production rates approaching equilibrium.

### Microscopy

All imaging was performed using an epifluorescence microscope custom-built on a Ti Eclipse body with PFS focus stabilization system (Nikon, Düsseldorf, Germany), built upon and modified from a previous version of the system as described in (Martens *et al*. 2024). The setup is equipped with five laser lines covering 405 nm, 488 nm, 638 nm, 750 nm (Oxxius, Lannion, France), and 561 nm (Novanta, Boston, USA) wavelengths which are coupled into a 70 μm diameter multi-mode fibre (CeramOptec, Bonn, Germany) by a protected silver reflective collimator (RC04FC-P01, Thorlabs), and then expanded and focused by several optical elements and a dichroic ZT405/488/561rpc (Chroma, Bellows Falls, VT, USA) onto the back focal plane of a CFI Apo TIRF 60 x oil objective (NA 1.49, Nikon). To ensure uniform illumination and eliminate speckle patterns, the fiber is mechanically vibrated using a small eccentric mass motor. The heating stage was set to 37°C for all imaging experiments. Fluorescence was recorded by a Prime BSI sCMOS camera (Teledyne Photometrics, Tucson, AZ, USA; 107 nm pixel size) and using ZET405/488/561m-TRF (Chroma) and bandpass filters ET510/80m (green Dendra2Hfx) and ET610/75m (orange Dendra2Hfx). The microscope setup was controlled by μManager 2.0 (Edelstein *et al*. 2014) combined with Pycro-Manager (Pinkard *et al*. 2021), while laser triggering was managed by a TriggerScope 4 (Advanced Research Consulting, Newcastle, CA, USA).

### Agarose pads

Agarose pads were prepared with 1.5 (w/v%) low-melting (Sigma-Aldrich, Germany) agarose and the respective HV-Cab medium contained the needed selection additives. Microscopy slides and coverslips were cleaned in 1 M KOH overnight. The agarose was melted at 70 °C for 20 min and cooled to 42 °C once fully melted. The melted agarose was then spotted onto dimpled microscopy slides (Thermofisher, Germany) and covered with coverslips to solidify.

For sptPALM imaging, plane microscopy slides and coverslips were subsequently cleaned with detergent, dH_2_O, 70% Ethanol and 1 M KOH. Gene Frame stickers (Thermofisher, Germany) were used to spot the samples onto the microscopy slides and then covered by cover slips.

### Cell morphology imaging and analysis

To analyze H119 and WR806 cell morphology, the respective strains were inoculated in HV-Cab or HV-YPC medium until stationary phase over an OD_600_ of 2 was reached. All cultures were then sub-cultured to an OD of 0.05 and grown until they reached OD 0.1. Cultures were prepared for imaging by centrifuging 2 mL of culture at 4000 rpm for 5 min and washed with the respective medium. Cell pellets were then resuspended in medium and spotted onto agarose pads. Brightfield images were obtained by acquisitions of brightfield movies (500 frames, 50 ms) and averaging the images using Fiji 1.54f (Schindelin *et al*. 2012). Cells were segmented using StarDist 0.3.0 (Schmidt *et al*. 2018; Weigert *et al*. 2020) with a pretrained model for *H. volcanii*, and cell shape descriptors (length, width, area, aspect ratio (length/width), circularity (4π × area/perimeter²)) were measured. Wrongly segmented cells were manually corrected. For significance testing, a Mann-Whitney-U test was performed.

### Diffraction-limited fluorescence imaging and data analysis

Cell samples were grown to stationary phase in HV-Cab medium prior to imaging and sub-cultured with the specified conditions into fresh HV-Cab with 0, 0.25, or 4 mM tryptophan and grown to the stated time points. Prior to imaging, cells were harvested in a microcentrifuge at 4000 rpm for 5 min and washed two times with fresh medium. Cells were then resuspended in a smaller volume of fresh medium and spotted onto agarose pads. Prior to fluorescence read-out, brightfield images were obtained to serve for cell segmentation. Green Dendra2Hfx (Turkowyd *et al*. 2020) was excited with a 488 nm laser at a laser power of 2 W/cm^2^ for 500 frames with 50 ms exposure. To obtain average fluorescence intensities per cell area, images were analyzed using a custom Python script. First, brightfield images were averaged and segmented with StarDist 0.8.5 (Schmidt *et al*. 2018; Weigert *et al*. 2020) with a pre-trained model for *H. volcanii*. Cells that were not properly segmented or cut-off by an image edge were manually excluded. The average fluorescence intensity per cell was calculated from averaged fluorescence images, normalized to the background intensity. To quantify the number of cells carrying fluorescence spots, cells were counted manually.

### UV light exposure of cell cultures for sptPALM imaging

WR806 pUE001-Cas1:Dendra2Hfx was grown from a single colony to stationary phase in HV-Cab medium supplied with thymidine and uracil (without tryptophan). Prior to imaging, the culture was sub-cultured to OD_600_ 0.02 with a final concentration of 0.25 mM tryptophan and grown overnight under inducing conditions. On the next morning, the culture was diluted to an OD_600_ 0.1. To expose the cells to UV-light, 3 mL of the culture were centrifuged in a 48-well plate. Before UV light exposure using a Thorlabs M265L5 UV lamp (265 nm, 38.4 mW, 440 mA), the supernatant was briefly removed to prevent UV absorption by the medium. Cells were exposed to the UV light for 235 ms (50 J/m^2^), 470 ms (100 J/m^2^) and 940 ms (200 J/m^2^), respectively. Afterwards, cells were quickly resuspended in 1 mL HV-Cab medium with uracil, thymidine and 0.25 mM tryptophan and grown for varying recovery times at 42 °C and 180 rpm. Before imaging, cells were centrifuged at 4000 rpm for 5 min and resuspended in a smaller volume of medium, before spotting onto agarose pads. Cell samples with the shortest recovery time of 15 min were spotted directly onto prepared agarose pads.

### sptPALM imaging and analysis

First, brightfield images (200 frames, 50 ms exposure) were obtained to record cell outlines. To track single Cas1:Dendra2Hfx fusion proteins, primed photoconversion to photoconvert Dendra2Hfx using a 488 nm and 750 nm laser was used (Turkowyd *et al*. 2017). At least 10000 frames with an exposure time of 20 ms each were recorded for each field of view. To control FP activation, the 488 nm laser was strobed for 1 ms at the beginning of each frame and laser power was increased continuously during the acquisition. Converted Dendra2Hfx was recorded by the 561 nm laser, strobed for 10 ms in the middle of each frame to avoid motion blur. The first 1000 frames of each acquisition were exposed only to the 561 nm laser to bleach noise and discarded from analysis.

Localization finding and single-particle tracking was performed with constant parameters for all datasets to ensure comparability (Supplementary Table S4, 5). Localization of emitters was performed using ThunderSTORM 1.3 (Ovesný *et al*. 2014). Tracking was performed by custom software (*swift* v0.4.3, Endesfelder lab, beta-test software available upon request to the authors) and mean jump distances (MJD) and motion types were analyzed. A Mann-Whitney-U test was performed to investigate differences between conditions. U-values were normalized to the respective sample size of both compared groups. The untreated control data set was randomly split into two parts to serve as a control condition.

### Software

Custom scripts were written with Python 3.11.5 using pandas, matplotlib, numpy, skimage, csbdeep, and tiffile libraries. GraphPad Prism 10.4.1 (GraphPad Software, LLC) was used for plotting and statistical tests. Affinity Designer 1 (Serif, Ltd) was used to finalize figures. *swift* v0.4.3 (Endesfelder lab, beta-test software available upon request to the authors), Pycro-Manager (Pinkard *et al*. 2021) and Fiji (Schindelin *et al*. 2012) with plugins for micromanger (Edelstein *et al*. 2014), Stardist (Schmidt *et al*. 2018; Weigert *et al*. 2020) and Thunderstorm (Ovesný *et al*. 2014) were used for image analysis and tracking. FTIR spectra were analyzed using OPUS (Bruker, USA).

## RESULTS and DISCUSSION

### Characterization of the carotenoid-deficient WR806 strain

Natural carotenoid synthesis of *H. volcanii* is facilitated by the phytoene dehydrogenase encoded by *crtI* (HVO_2528). The produced carotenoids cause significant cellular autofluorescence under fluorescence excitation, a major caveat for SMLM imaging as it hinders the detection of single molecules (Turkowyd *et al*. 2020). To overcome this limitation, the “albino” strain WR806 (DS70-wildtype (ΔpHV2), *ΔpyrE2*, *ΔtrpA*, *ΔleuB*, *ΔhdrB*, *ΔcrtI*), with disrupted carotenoid biosynthesis by a *ΔcrtI* deletion was created (Turkowyd et al., 2020). Given the unresolved open question of whether carotenoid depletion might affect native cellular processes, particularly membrane biophysics, we now performed detailed comparative analyses between WR806 and the red carotenoid-producing strain H119 (DS70-wildtype (ΔpHV2), *ΔpyrE2*, *ΔtrpA*, *ΔleuB*; (Allers *et al*. 2004)). Analysis of lipid extracts from H119 and WR806 cultures via visual inspection and absorption spectroscopy confirmed absence of carotenoids in WR806 extracts (Figure 1A). To characterize the molecular composition of these extracts, we performed Fourier-transformed infrared (FTIR) spectroscopy. The FTIR spectra, normalized to intensity of the asymmetric CH_2_ vibration peak at 2925 cm^-1^ associated with the lipid side chains (Movasaghi, Rehman and Rehman 2008), demonstrated remarkable compositional similarity between WR806 and H119 lipid extracts (Figure 1B). While absolute peak intensities varied due to different lipid concentrations, second derivative analysis confirmed the nearly identical chemical composition of both extracts. Notably, the characteristic bacterioruberin peak at 1650 cm^-1^ (Noby, Khattab and Soliman 2023) showed only minimal variation between samples.

We assessed the biophysical membrane properties of the lipid extracts using Langmuir-Blodgett monolayer compression assays under two conditions: physiological (pH 7.5, 42°C; Figure 1C) and SMLM imaging conditions (pH 7.5, 37°C; Supplementary Figure S2A). The surface-pressure-area isotherms, normalized to lipid molecule numbers quantified by a phosphate assay, were analyzed for changes in compressibility — a sensitive indicator of phase transitions and alterations in lipid ordering (Risović, Frka and Kozarac 2011). The identical slopes observed under both conditions, with no inflection points and thus no evidence of phase transitions, suggested equivalent biophysical behavior between the extracts. The observed lateral displacement of isotherms along the x-axis can be attributed to quantification limitations of the phosphate assay, which fails to detect non-phosphate-containing membrane components such as sterols, quinones, and carotenoids. Given that *H. volcanii* membranes contain significant amounts of apolar lipids (Kellermann *et al*. 2016), this technical limitation affects the accuracy of absolute lipid concentration measurements. Analysis of isothermal compressibility (Supplementary Figure S2C, D) revealed similar values for both lipid monolayers at surface pressures between 5-20 mN/m. Above 20 mN/m, H119 extracts exhibited slightly higher compressibility, differing by approximately 10 mN/m from WR806 extracts. Together, the monotonic nature of the isotherm curves at both 37°C and 42°C, combined with overlapping compressibility modulus values, demonstrates that both H119 and WR806 strains maintain similar membrane mechanical properties. This biophysical equivalence suggests that membrane properties should not differentially affect cell biology between the strains. Notably, these findings align with a recent study on carotenoid-deficient mutants of the gram-negative *Methylobacterium extorquens*, which similarly exhibited no significant changes in growth or membrane biophysical phenotype (Rizk *et al*. 2021).

We next examined living cells cultured at 42°C to assess growth dynamics and morphological characteristics. Growth analysis revealed comparable doubling times between H119 and WR806 strains in HV-Cab medium (4.6 ± 0.9 hrs and 4.5 ± 0.9 hrs, respectively; Figure 1D). In HV-YPC medium, doubling times showed minor variation but remained within one standard deviation (H119: 3.9 ± 0.5 hrs; WR806: 3.2 ± 0.5 hrs; Figure 1D). When transferring cultures between media types, adaptation occurred more rapidly during the transition from HV-Cab to HV-YPC (Supplementary Figure S2B), with strain-specific variations remaining within one standard deviation.

Morphological analysis revealed that while cell circularity and aspect ratio (AR) remained consistent between strains, the cell area exhibited significant differences in both media conditions. WR806 cells were consistently larger, with median areas of 3.9 ± 1.5 µm² (HV-Cab) and 4.1 ± 1.5 µm² (HV-YPC), compared to H119 cells with 3.6 ± 1.2 µm² (HV-Cab) and 3.9 ± 1.4 µm² (HV-YPC) (P < 0.0001 for HV-Cab; P = 0.0002 for HV-YPC). At this point we can only speculate and may attribute this size difference to the absence of carotenoids: while our *in vitro* experiments showed membrane compressibility remains unaffected, the lack of carotenoids might influence cell volume expansion. Previous studies have found that lipidome minimalization resulted in increased cell volumes in *Mycoplasma mycoides* minimal cells, highlighting the possible effects of depleted membrane components on cell area (Justice et al., 2024). Importantly, the preserved cellular length/width ratios and circularity metrics indicate intact cell shape regulation, which is crucial for pleomorphic *H. volcanii* (Duggin et al., 2015; Patro et al., 2023).

Collectively, these results validate WR806 as an imaging strain that maintains an almost native-like biophysical and physiological properties while enabling single-molecule detection capabilities.

### Increased control over protein expression using a novel plasmid system

Derivatives of pTA231 are often used in *H. volcanii*, which feature the constitutive promoter p.syn for POI expression and the *trpA* selection marker under p.fdx control (Allers *et al*. 2004; Berkemer *et al*. 2020). Another plasmid type are derivatives of the plasmid pTA962, incorporating the tryptophan-inducible promoter p.tna and the selection markers *hdrB* and *pyrE2* (Allers *et al*. 2010). Inducible promoter systems offer significant advantages over constitutive ones, primarily by allowing precise control over expression levels. This control is particularly crucial for cell-toxic proteins, highly regulated proteins, or highly stable proteins. However, the pTA962 system in WR806 strain presents a limitation: Because the *trpA* deletion in WR806 disrupts internal tryptophan synthesis, cells must be continuously supplied with external tryptophan (0.25 mM) for normal growth. This couples selection and induction, as the same molecule serves both functions, resulting in constant induction of protein expression under standard growth conditions. This coupling effectively negates the benefits of an inducible system. To address this limitation, we developed a novel plasmid: pUE001 encodes the *trpA* gene, enabling internal tryptophan synthesis. Combined with p.tna for inducible POI expression, this decouples selection from induction, restoring the functionality of the inducible system while maintaining the advantages of controlled protein expression in *H. volcanii.* pUE001 was constructed by cloning the p.tna promoter from pTA962 into the backbone of pTA231, resulting in a plasmid with the *trpA* selection marker under the constitutive promoter p.fdx and a tryptophan inducible p.tna promoter for POI expression. The plasmid was further engineered to enable labeling of POIs with a SMLM-compatible FP by inserting the coding sequence for Dendra2Hfx (Turkowyd *et al*. 2020) downstream of the promoter (Supplementary Figure S1, Addgene # 234669).

To characterize the novel pUE001 expression system in biological applications we also constructed versions that encode FtsZ1:Dendra2hfx and Cas1:Dendra2Hfx fusion proteins. These proteins were selected based on our previous challenges (data unpublished) encountered with the pTA962 expression system: The highly stable protein FtsZ1 was chosen due to its tendency to form protein aggregates and inclusion bodies which make it an ideal candidate to test for improved expression control. Cas1 was selected as a naturally low-abundant and seemingly highly regulated protein, as it had proven difficult to express in +previous pTA962 fusion constructs.

First, we conducted quantitative fluorescence and OD_600_ measurements of WR806 strains carrying either pUE001-Dendra2Hfx or pUE001-FtsZ1:Dendra2Hfx, using plasmid-free WR806 as a negative control. Cas1 as a lowly expressed (Brendel *et al*. 2014; Jevtić *et al*. 2019) was too low abundant to be reliably measured in plate reader experiments. To characterize induction levels and fluorescence response over time, we grew each strain in HV-Cab medium with varying tryptophan concentrations (0-5 mM for WR806 carrying pUE001; 0.25-5 mM for WR806 and WR806 carrying pTA962) for 70 hours. We normalized fluorescence values to optical density measurements (Figure 2; raw data in Supplementary Figure S3).

The normalized fluorescence data (Figure 2A, B, D, E) reveals distinct patterns corresponding to different growth phases. During lag phase, the fluorescent protein production rate (r_FP_) exceeds the cell division rate (r_D_), resulting in rapidly increasing fluorescence per cell. In exponential phase, r_FP_ and r_D_ reach equilibrium, manifesting as a plateau in the normalized fluorescence data. When entering stationary phase, r_FP_ again exceeds r_D_, leading to an increase of the fluorescence per cell.

Analysis of steady-state fluorescence intensities during exponential growth phase shows optimal induction at 3-4 mM tryptophan concentration (Figure 2 C, F), consistent with previous findings (Large *et al*. 2007). While pTA962-carrying strains exhibit higher absolute fluorescence intensities, pUE001-carrying strains demonstrate a more linear correlation between tryptophan concentration and expression levels. Maximum induction occurs at 3 mM for pTA962-carrying strains compared to 4 mM for pUE001-carrying strains. Crucially, since WR806 carrying pUE001 can grow without tryptophan supplementation, uninduced cultures show baseline fluorescence comparable to the empty WR806 control in exponential phase growth. This comparison of pTA962 and pUE001 steady-states during exponential growth reveals pTA962’s limited induction range when used in a strain without a chromosomal copy of *trpA* like WR806 at both high and low tryptophan concentrations, highlighting pUE001’s superior expression control and induction linearity.

This also underscores the importance of preculture conditions. All strains in these plate reader experiments were grown for about three days into stationary phase before sub-culturing, resulting in initial fluorescence intensities which are roughly equivalent to the 70-hour timepoint (marked by asterixis in Figure 2A, B, D, E). Since WR806 carrying pTA962 requires externally supplied tryptophan for growth, it shows higher initial fluorescence than WR806 carrying pUE001 strains (asterixis in Figure 2A, B, D, E). This poses a major limitation for ensemble studies by precluding true zero-expression conditions. In pUE001 0 mM tryptophan conditions, we observed increased fluorescence during stationary phase (Figure 2A, D), likely due to tryptophan accumulation in the media from e.g. degraded cell debris.

Finally, this plate reader dataset highlights protein stability as a crucial factor in expression level measurements. FtsZ1, a filament-forming and highly stabilized protein (Liao *et al*. 2021; Pende *et al*. 2021), likely has a longer lifetime than artificial, cytosolically expressed Dendra2Hfx. This is nicely reflected in higher initial values for FtsZ1:Dendra2Hfx-expressing strains (Figure 2D, E). While pUE001-Dendra2Hfx carrying strains show a fast decrease of fluorescence between stationary phase (mimicking preculture conditions) and experiment start (Figure 2A), this effect is not visible for pUE001-FtsZ1:Dendra2Hfx strains (Figure 2D). We attribute this to fast cytosolic Dendra2Hfx dilution and degradation during experimental setup (time between preparation of the experiment and start of the plate reader run ∼1 hour), while stabilized FtsZ1:Dendra2Hfx remains unaffected. This effect is also absent in by 0.25mM tryptophan continuously induced WR806 carrying pTA962 strains (Figure 2B, E) and will be readdressed further below.

### Decoupling selection and induction enables tight expression control and reveals potential promotor-independent regulation of *trpA*

Following ensemble studies, we performed single-cell sensitive diffraction-limited imaging of WR806 expressing three POIs: Cas1:Dendra2hfx, FtsZ1:Dendra2Hfx, and cytosolic Dendra2hfx, using both pUE001 and pTA962 expression systems. This approach enabled detection of the low-abundant Cas1 protein (Brendel *et al*. 2014; Jevtić *et al*. 2019). Notably, we found that plasmid-based expression of fusion proteins is not universally successful, as demonstrated by our attempts to express FtsZ1:Dendra2Hfx from pTA231 under the constitutive p.syn promoter (Supplementary Figure S4), although cytosolic fluorescent proteins expressed successfully (Turkowyd *et al*. 2020). Similar expression challenges have been reported elsewhere (Ithurbide *et al*. 2024). We have not investigated this further but recommend to check combinations of imaging strains and expression systems carefully for each POI and experimental set up.

We cultured strains with high (4 mM) and low (0.25 mM) tryptophan concentrations, with WR806 pUE001 strains additionally grown without tryptophan supplementation. Imaging after 18 hours in exponential phase demonstrated successful expression of all POIs using pUE001 and confirmed reliable p.tna promoter induction, independent of the plasmid-encoded selection marker *trpA* (Figure 3). This data reinforces our plate reader findings and extends them to Cas1:Dendra2Hfx expression. While pTA962 cannot be grown without constant p.tna promoter induction as seen by leaky expression across all POIs, pUE001’s decoupled induction resulted in tight expression control under tryptophan-free conditions (Figure 3). This feature allows for fine-tuned expression at tryptophan concentrations between 0-0.25 mM of highly regulated, toxic, or growth-inhibiting proteins, including CRISPR-Cas components (Figure 3A) and stabilized proteins like FtsZ1 (Figure 3C). Intriguingly, since *trpA* is under control of the constitutive p.fdx promoter, our results confirm our hypothesis of promoter-independent regulation, though we did not investigate this mechanism further.

**Figure 3.**
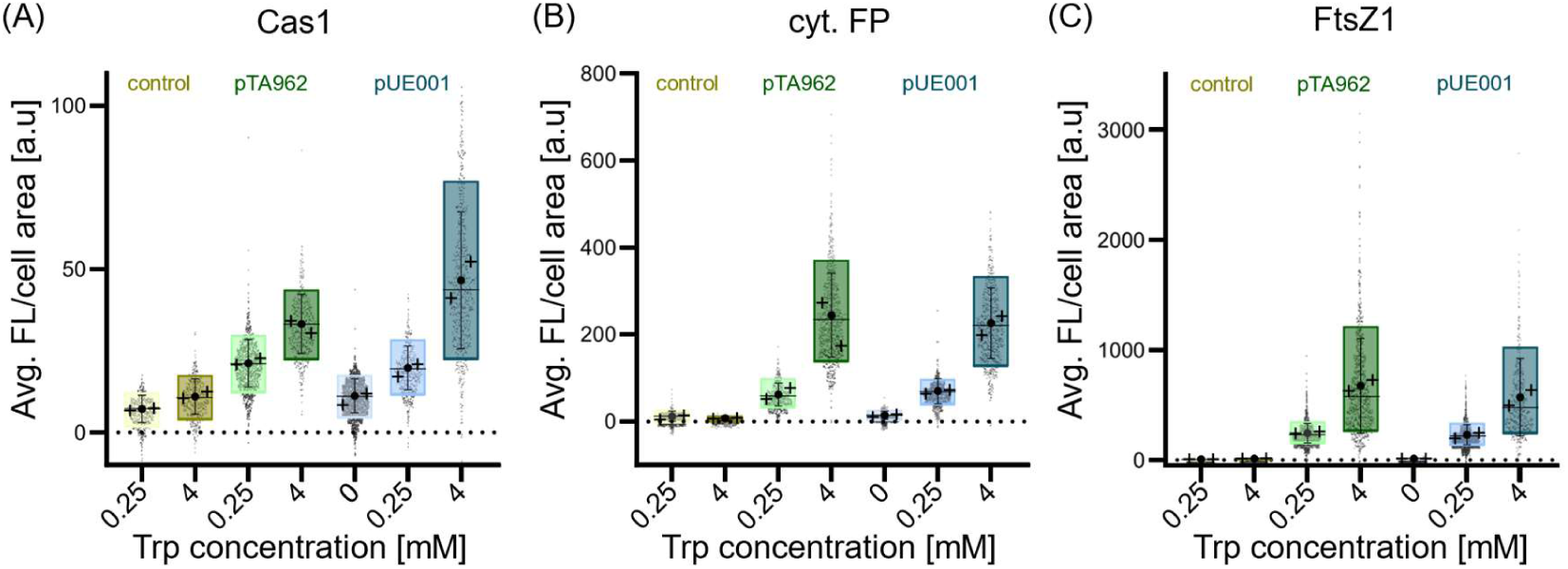
Single-cell analysis of Dendra2Hfx fluorescence expression in *H. volcanii* strain WR806. Quantification of background-corrected green fluorescence intensity per cell area using diffraction-limited fluorescence microscopy. Cells expressed different Dendra2Hfx constructs from either pTA962 (green) or pUE001 (blue) plasmids in three configurations: (A) Cas1:Dendra2Hfx fusion protein, (B) cytosolic Dendra2Hfx, and (C) FtsZ1:Dendra2Hfx fusion protein. Cells were inoculated at an OD600 of 0.1 and cultured under inducing conditions for 18 hrs, allowing them to re-enter exponential growth phase and reach steady-state equilibrium between fluorescence production and growth rate, as previously demonstrated in Figure 2. Plasmid-free WR806 (yellow) served as negative control. Data from biological duplicates (n > 1000 cells total) are shown as individual data points and summarized in box plots showing overall mean (circle), individual replicate means (crosses), median (horizontal line), 10th-90th percentile range (box), and standard deviation (whiskers).

This data further reveals protein-specific expression patterns, particularly evident in comparing average intensity levels between Cas1:Dendra2Hfx and FtsZ1:Dendra2Hfx expressing strains (Figure 3A, C). Furthermore, we can assume that cytosolic Dendra2Hfx appears to minimally interfere with cellular processes, making it a potential benchmark to estimate protein abundance in *H. volcanii*.

Notably, high tryptophan concentrations (4-5 mM) induced some autofluorescence in WR806 (Figure 3A), with average intensities increasing from 7 ± 4 a.u. in WR806 0.25 mM to 11 ± 5 a.u. in WR806 4 mM above background. This autofluorescence increase upon high tryptophan concentrations might pose a problem under SMLM conditions. However, high tryptophan concentrations of 4 mM are typically not used for imaging cell biology as severe POI overproduction would most likely show alterations. Furthermore, OD measurements of WR806 under high tryptophan concentrations show accelerated entry into stationary phase (Supplementary Figure S3). This effect might be related to indole, the primary product of tryptophan degradation by *tnaA*-encoded tryptophanase as known for gram-negative bacteria and halophilic archaea (Boya *et al*. 2021). Indole has been shown to function as a signaling molecule affecting biofilm formation, cell division, and virulence in prokaryotes (Di Martino *et al*. 2003; Lee and Lee 2010), including persister cell formation in *H. volcanii* (Megaw and Gilmore 2017). Studies in *E. coli* have shown indole’s interaction with the stationary phase-inducing sigma factor δ^S^ (Wang, Ding and Rather 2001; Lee and Lee 2010; Joffré *et al*. 2022) and transient spikes in intracellular indole concentration during exponential-to-stationary phase transition (Gaimster *et al*. 2014). Similar mechanisms might exist in *H. volcanii*. While the global effect of indole accumulation on cellular processes may explain cellular stress and thus, next to increased fluorescence background by aromatic tryptophan itself, might contribute to autofluorescence at high tryptophan concentrations, the overall increase in autofluorescence in our hands remains negligible when imaging of fluorescent protein-expressing cells. Comparing the autofluorescence levels of WR806 with signal of low-abundant proteins like Cas1 at 4mM tryptophan concentrations, the FP signal still show an about four-times increased signal with a mean fluorescence intensity of 47 ± 21 a.u.. For FtsZ1 this difference is even more pronounced, as the signal is about 50 times higher with a mean fluorescence intensity of 571 ± 351 a.u. (Figure 3).

### Tryptophan synthesis by *trpA* can prevent inclusion body formation in WR806 carrying pUE001 expression constructs

Conducting the single-cell diffraction-limited imaging as shown in Figure 3, indicated that handling of precultures and growing conditions critically affects imaging outcomes. We frequently observed fluorescent foci and higher cellular fluorescence intensities WR806 strains expressing FtsZ1:Dendra2Hfx (Figure 4A, B), even though the cultures were regrown for 18h after sub-culturing from stationary phase precultures. To further investigate the accumulation of fluorescence during stationary phase (Figure 2) and formation of fluorescent foci (Figure 4A,B), we quantified the number of fluorescent spots per cell over time (Figure 4C). We used minimal tryptophan concentrations for each strain: 0 mM for pUE001-FtsZ1:Dendra2Hfx and 0.25 mM for pTA962-FtsZ1:Dendra2Hfx and re-inoculated the strains from three-days stationary phase pre-cultures. Fluorescent spots were counted at 0, 6, 12, and 18 hours. Cultures directly from stationary phase precultures at 0h showed similar proportions of cells with fluorescent spots: 75 ± 6% for pTA962 and 77.8 ± 2% for pUE001 strains (Figure 4C). Upon sub-culturing into fresh medium, we observed distinct temporal patterns: WR806 carrying pUE001 showed complete elimination of fluorescent foci after 12 hours, while WR806 carrying pTA962 maintained a minimum of 45 ± 13% cells with spots even after 18 hours (Figure 4C), the condition which was measured in Figure 3. This decrease of cells exhibiting spots correlated with the declining average cellular fluorescence intensities (Figure 4B). Notably, spot characteristics differed between strains: WR806 carrying pTA962 typically showed single, larger, and brighter foci per cell, while WR806 carrying pUE001 exhibited multiple smaller, less intense spots (Figure 4A, B).

**Figure 4.**
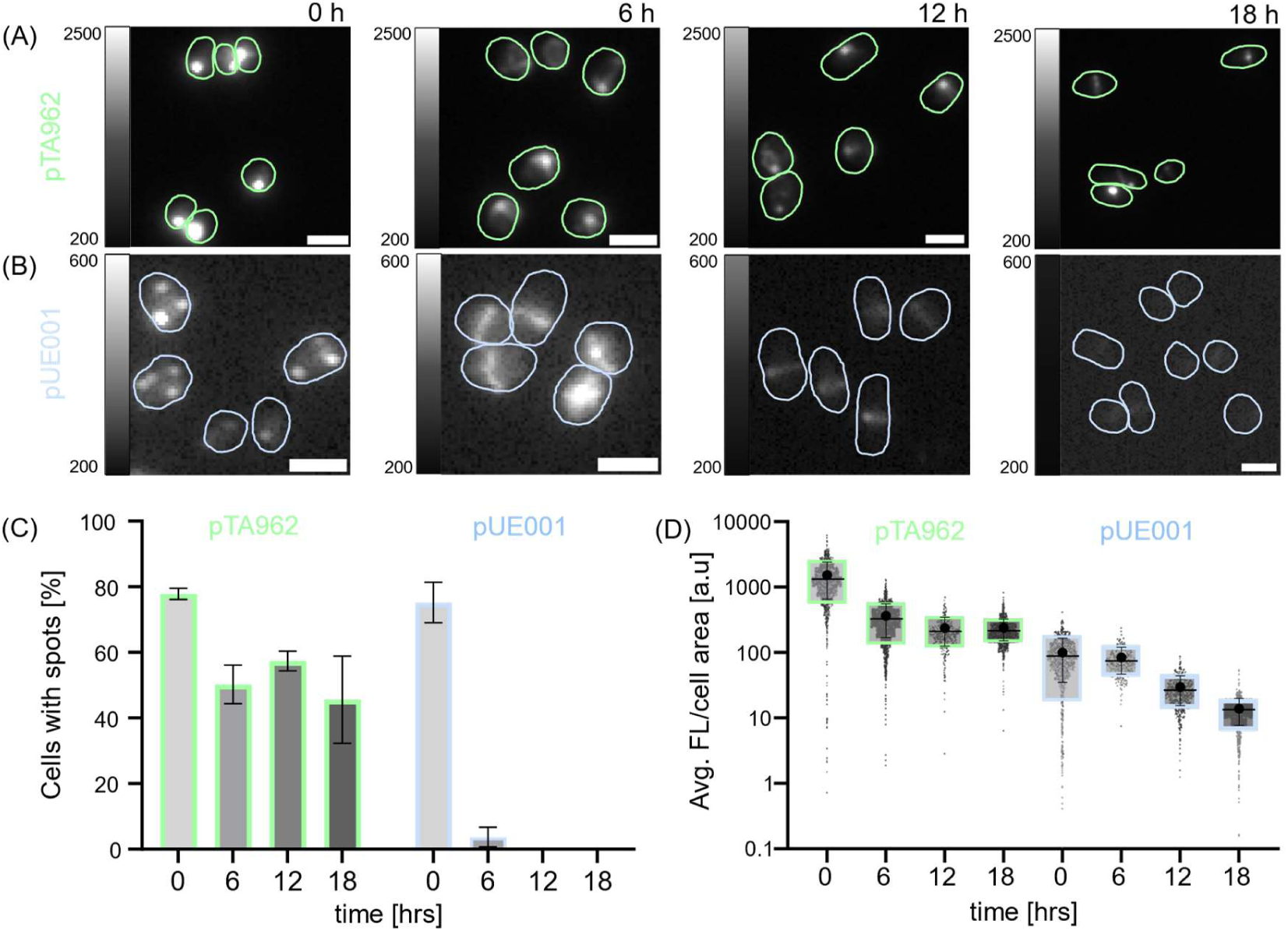
Time-course analysis of FtsZ1-Dendra2Hfx localization and expression in *H. volcanii* strain WR806. Cells were taken from a three-day stationary pre-culture supplied with 0.25 mM tryptophan in the case of WR806 pTA962 and 0 mM tryptophan in the case of WR806 pUE001, inoculated under the same conditions and imaged after 0, 6, 12 and 18 hours by diffraction-limited epifluorescence microscopy (18 hrs samples are the same dataset as in Figure 3). Measurements were taken in biological duplicates. (A, B) Representative fluorescence images showing green Dendra2Hfx fluorescence signal in quantitative grayscale intensities. Scale bar 2 µm. (C) Mean fluorescence intensity per cell area plotted on logarithmic scale. (D) Percentage of cells displaying fluorescent spots (in addition to FtsZ1 ring structures).

Given their brightness and persistence we thus speculate that foci in WR806 carrying pTA962 represent inclusion bodies (IBs) and result from constitutive protein expression under normal growth conditions. IBs, typically associated with protein overproduction and cellular stress (Fahnert, Lilie and Neubauer 2004; Landgraf *et al*. 2012; Kopp *et al*. 2023), pose significant challenges for both biotechnology applications and fluorescence imaging by deviating from native cell biology and compromising the applications. In contrast, the transient spots in pUE001 likely might represent smaller FtsZ1 oligomers. During stationary phase, when cell division is arrested, overexpressed FtsZ1 might form these filaments, which subsequently could be incorporated into functional FtsZ1 rings upon resumption of cell growth and cell division. This process indicates a FtsZ1:Dendra2Hfx redistribution from small spots to distinct mid-cellular rings between 0 and 6 hours after restarting growth, where average fluorescence intensity is maintained but reorganized protein localization shows a clear shift (Figure 4D). As cultures enter exponential phase, per-cell fluorescence intensity decreases, consistent with our plate reader data (Figure 2B). The remarkable stability of FtsZ1 is further demonstrated by the detection of faint FtsZ rings even after 18 hours without induction in WR806 pUE001-FtsZ1:Dendra2Hfx, likely due to stochastic p.tna promoter activity (Figure 4D).

We also observed fluorescent spots, albeit at much lower frequencies, in strains expressing Cas1:Dendra2Hfx (0.75% of cells for pTA962 and 0.15% for pUE001, data not shown), while strains expressing cytosolic Dendra2Hfx never showed foci. This protein-specific aggregation pattern suggests a strong correlation between protein function and aggregation propensity, while confirming Dendra2Hfx as an ideal, truly monomeric FP for *H. volcanii* studies due to its minimal interference with cellular processes.

These findings emphasize the importance of careful strain, expression system, and induction parameter selection based on the POI. Furthermore, they demonstrate the superior performance of the pUE001 system in terms of rapid adaptation to non-inducing conditions and overall experimental utility.

### Cas1 might be involved in UV-light associated repair of DNA damage

CRISPR-Cas systems, initially hypothesized to function in DNA repair before their role in adaptive immunity was discovered (Makarova *et al*. 2002), are increasingly recognized for their diverse cellular functions. Recent studies have revealed strong interconnections between Cas proteins and fundamental cellular processes, including gene regulation, virulence, signal transduction, and DNA repair mechanisms (Faure, Makarova and Koonin 2019). The adaptation complex protein Cas1, with its nuclease activity, demonstrates particular utility in DNA repair through its ability to cleave branched DNA substrates. This activity has been documented across domains, from *E. coli* to archaeal *H. volcanii*, suggesting a conserved link between DNA repair and Cas1 function (Babu *et al*. 2011; Rollie *et al*. 2015; Wörtz *et al*. 2022). The direct interaction between Cas1 and nucleotide excision repair (NER) proteins associated with UV damage repair in *H. volcanii* (Wörtz *et al*. 2022), coupled with an increased UV sensitivity in *E. coli* Cas1 deletion strains (Babu *et al*. 2011), prompted us to investigate Cas1’s role in UV-induced DNA repair using single-particle tracking photoactivated localization microscopy (sptPALM).

Our novel pUE001 plasmid, which demonstrates superior performance compared to pTA962 in WR806, enables precise control of expression levels between 0 and 0.25 mM tryptophan — a crucial advancement for sensitive sptPALM measurements. We chose to express pUE001 Cas1:Dendra2Hfx in WR806 also encoding for the native, untagged genomic Cas1, as studies indicate that expressing tagged POIs in their respective deletion strains can significantly disrupt cellular function (Duggin *et al*. 2015; Liao *et al*. 2021). To ensure cells were in equilibrium state, where r_FP_ and r_D_ are equal (Figure 2), cultures were kept in early exponential phase preparatory to imaging. Given the low abundance and tight regulation of Cas proteins (Brendel *et al*. 2014), this approach, combined with minimal induction (0.25 mM tryptophan), ensured the required low POI expression and thus native-like functional behavior of defense and possible DNA repair mechanisms.

In *in vivo* single-molecule studies of CRISPR-Cas interference, Type-II Cas9 (Knight *et al*. 2015; Jones *et al*. 2017; Martens *et al*. 2019) and Type-I Cascade complexes (Turkowyd *et al*. 2019; Vink *et al*. 2020) both exhibit rapid diffusion and transient binding while screening DNA for protospacer adjacent motif (PAM) sites. Similar dynamics can be assumed for Cas1 in *H. volcanii*, as the protein takes over a major function in the PAM verification process during adaptation (Nuñez *et al*. 2014; Wang *et al*. 2015). Once Cas1 becomes actively involved in e.g. DNA defense or DNA repair processes, a significant slowdown of this fast-diffusive signal is to be expected.

To investigate this, we induced cyclobutane pyrimidine dimers (CPDs) — the predominant form of UV-associated DNA damage—using low (50 J/m²), medium (100 J/m²), and high (200 J/m²) doses of 265 nm UV light, based on previous studies of the UV damage-associated NER uvrABCD system in *H. volcanii* (Lestini, Duan and Allers 2010) and followed the molecular dynamics at 15, 60, and 180 minutes post-exposure.

Analysis of weighted mean jump distances (wMJD) revealed significant UV damage-induced slowdown of a mobile fraction of Cas1 (P < 0.0001) across all conditions (Figure 5 and Supplementary Figure S5). Normalized U-values demonstrate both time-and dose-dependent responses, with lower U-values indicating stronger effects. Under low UV exposure, we observed a gradual mobility decrease, with U-values declining from 0.47 (15 min) to 0.45 (180 min). The most pronounced response occurred with medium exposure (100 J/m²) after 180 minutes (U-value: 0.34). Interestingly, high UV doses (200 J/m²) produced an initial slowdown at 15 and 60 minutes followed by a shift back to match more closely again to the original distribution at 180 minutes. Cytosolic Dendra2Hfx as a control did not show any change in diffusive behavior (data not shown).

**Figure 5.**
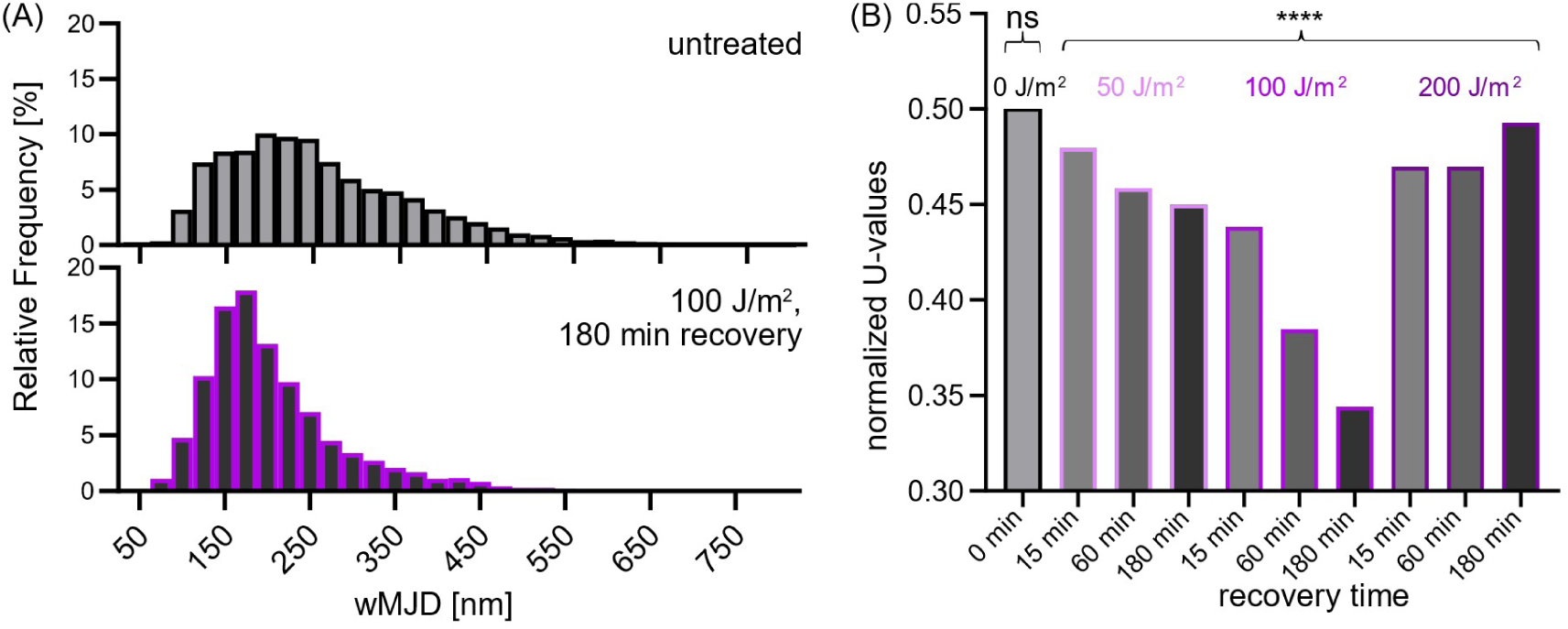
Analysis of Cas1-Dendra2Hfx single-molecule dynamics in response to UV-light damage in *H. volcanii* strain WR806 pUE001 as measured by single-particle tracking microscopy. Exponentially growing cells cultured without tryptophan were sub-cultured with 0.25 mM tryptophan for inducing Cas1 protein expression overnight. Cells were reinoculated under the same conditions to OD 0.1 and after 2 hours of growth, exposed to UV-light radiation at 265 nm at 0, 50, 100, or 200 J/m² and measured after a recovery time of 0, 15, 60, and 180 minutes post-exposure. (A) Distribution of Cas1 dynamics comparing untreated cells with cells showing maximal response (180 minutes recovery after 100 J/m² UV exposure). Dynamics are quantified by weighted mean jump distances (wMJD) from individual single-molecule trajectories across two biological replicates. (B) Statistical comparison of all experimental conditions with the untreated control using the Mann-Whitney-U-test and visualized by normalized U-values to account for difference in sample sizes (all wMJD distributions are provided in Supplementary Figure S5). Furthermore, the untreated data was split into two halves and tested against each other as a control. ns indicates not significant, **** indicates P < 0.0001.

Our experiments did not directly measure interactions between Cas1 and DNA repair pathways, so we can only speculate about mechanisms explaining the observed dynamics. The timing of Cas1 slowdown differs from *E. coli*’s rapid NER response, where uvrAB proteins react within 5-15 minutes of UV damage (Stracy *et al*. 2016). However, with regard to the comparably slow generation turnover of *H. volcanii*, it seems plausible that a UV damage response might occur later if e.g. coupled to transcription processes (Savery 2007; Pérez-Arnaiz *et al*. 2020). Interpretation of Cas1 dynamics at 200 J/m² requires caution, as this dose exceeds previously reported UV exposures for *H. volcanii* (maximum 100 J/m²; (Lestini, Duan and Allers 2010; Wörtz *et al*. 2022)) and may trigger cell death. Additionally, other DNA damage repair systems, e.g. for double-strand-repair (Pérez-Arnaiz *et al*. 2020), might be differently involved and/or prioritized for different, i.e. higher UV dosages. Double-strand breakages are a more severe form of DNA damage when compared to CPDs and are known to result from strong UVC light (Rolfsmeier, Laughery and Haseltine 2010). Thus, NER-associated slowdown of Cas1 might occur later and/or less prominent. While technical constraints prevented observations earlier than 15 minutes post-UV-exposure, the gradual response patterns at lower doses suggest maximum Cas1 slowdown may occur beyond 180 minutes in these conditions.

### Conclusions

The study of molecular cell biology through fluorescence imaging techniques as a direct, visual approach fundamentally depends on expression systems and selection markers that minimally perturb native cellular processes. This requirement becomes particularly detectable in single-molecule imaging, where even subtle alterations can significantly impact observations. Given the still limited availability of established expression systems and selection markers in the archaeal model organism *H. volcanii*, we developed and characterized a novel expression system optimized for the autofluorescence-free imaging strain WR806.

Our work validates that the combination of the carotenoid-free strain WR806 and the novel pUE001 expression plasmid is an excellent tool for quantitative and low-induction fluorescence studies of live *H. volcanii* cells as the biophysical and physiological parameters remain unaffected by carotenoid depletion upon deletion of *crtI* and protein expression from pUE001 prevents artefacts such as inclusion body formation, and enables controlled and finetuned expression. We demonstrate through both ensemble and single-cell fluorescence measurements that decoupling induction from selection is vital to obtain robust data which closely reflects native cell biology. Our system enables precise control of protein expression levels for diverse proteins of interest — Cas1:Dendra2Hfx, FtsZ1:Dendra2Hfx, and cytosolic Dendra2Hfx — with fine-tunable regulation through tryptophan concentrations. This tight control represents a significant advancement over the previously used pTA962 system. Furthermore, our system offers enhanced practical utility, as expression levels can be effectively reset through an about 18h long sub-culturing of precultures.

Our findings establish that providing a copy of *trpA* is essential for proper control of the tryptophan-inducible p.tna promoter. More broadly, our work emphasizes the critical importance of careful consideration when combining selection markers, promoter systems, and strains, particularly in *H. volcanii*, which exhibits complex and sensitive pleomorphic characteristics (Patro *et al*. 2023). This strategic approach to expression system design proves especially crucial for maintaining near-native cellular conditions in molecular imaging studies.

We then investigated the interaction between native CRISPR-Cas Type I-B systems and DNA repair in *H. volcanii*. We observed a significant, dose- and time-dependent reduction in Cas1 protein mobility following UV-light associated DNA damage, with maximum response at 100 J/m² after 180 minutes recovery, suggesting interplay between UV damage repair mechanisms and CRISPR-Cas proteins. The involvement of Cas proteins in DNA repair has been documented across diverse organisms (Faure, Makarova and Koonin 2019). Cas1 specifically demonstrates nuclease activity on branched DNA substrates, particularly 5’-flaps, as shown *in vitro* using purified proteins from *E. coli* and *Sulfolobus solfataricus* (Babu *et al*. 2011; Rollie *et al*. 2015). This activity on structures commonly associated with DNA repair intermediates suggests a role for Cas1 in DNA maintenance. In *H. volcanii*, Cas1 exhibits functional similarity to the flap exonuclease Fen1 and associates with several replication and repair proteins, including the double-strand break repair protein Rad50, DNA mismatch repair MutS, and DNA helicase MCM (Wörtz *et al*. 2022).

Particularly relevant to our findings, Cas1 co-purifies with nucleotide excision repair (NER) proteins uvrA, uvrB, and uvrD (Wörtz *et al*. 2022), which are known mediators of UV-induced DNA damage repair (Lestini, Duan and Allers 2010). This association aligns with observations in *E. coli*, where *cas1* deletion increased UV sensitivity. Notably, double mutants of Cas1 and uvrABC homologues *ruvA*, *ruvB*, and *ruvC* show no additional UV sensitivity increase, suggesting these proteins operate in a shared pathway (Babu *et al*. 2011). Wörtz et al. (2022) proposed that recruitment of Cas1 to DNA repair complexes at damage sites might prevent Cas1-Cas2 adaptation complex formation, thereby inhibiting auto-immunity and positioning Cas1 as an accessory protein in DNA damage repair — a hypothesis supported by our observations.

Our findings reinforce the emerging view of DNA repair, replication, and defense as interconnected, dynamic networks, which highlights the expanded possible roles for CRISPR-Cas I-B systems beyond immunity. This work provides evidence for Cas1’s involvement in UV-light induced DNA damage response through SPT data, complementing previous biochemical and genetic studies.

## RESOURCES

All pUE001 plasmids are available on Addgene: #234669 (Dendra2Hfx), #234670 (Cas1:Dendra2Hfx), #234671 (FtsZ1:Dendra2Hfx).

## DATA AVAILABILITY

All data underlying the figures and supplementary figures is available under ZENODO (10.5281/zenodo.14845238).

## AUTHOR CONTRIBUTIONS

Conceptualization of the study by KJAM and UE. UE supervised the study with the help of AM, KJAM and JS. JW and PS created plasmids and strains. UA conducted and analyzed *in vitro* studies of lipid extracts. PS conducted and analyzed all other experiments. PS and UE designed figures and wrote the manuscript with the help of all authors.

## ACKNOWLEDGEMENTS

We thank Marlene Hecker for her help and expertise when reviving the FTIR setup, and Bartosz Turkowyd and Luis Gaiser for conceptualizing and establishing sample preparation protocols for single-particle tracking experiments investigating UV-light induced damage to cell biology. We further would like to acknowledge the constructive feedback from all members of the UE and JS laboratories during group discussions. We thank Titouan Bouvier D’Yvoire for critical reading of the manuscript.

## FUNDING

This work was financially supported by funding by the DFG (En1171/1-2 and Ma1538/25-2) in the frame of the DFG priority program SPP2141, by start-up funds (UE) and a stipend (UA) from Bonn University and as part of the Excellence Strategy of the federal and state governments of the excellence initiative by STEP funds (UE) and an Argelander Starter Kit (KJAM) from Bonn University; a stipend from the Alexander von Humboldt Foundation (KJAM); and DFG #527181518 to JS.

## CONFLICT OF INTEREST

The authors declare that there are no conflicts of interest.

## Notes

https://zenodo.org/records/14845238

https://www.addgene.org/Ulrike_Endesfelder/

### Competing Interest Statement

The authors have declared no competing interest.

## REFERENCES

1. Allers T, Barak S, Liddell S et al. Improved Strains and Plasmid Vectors for Conditional Overexpression of His-Tagged Proteins in *Haloferax volcanii*. Appl Environ Microbiol 2010;76, DOI: 10.1128/AEM.02670-09.

2. Allers T, Ngo HP, Mevarech M et al. Development of Additional Selectable Markers for the Halophilic Archaeon Haloferax volcanii Based on the leuB and trpA Genes. Appl Environ Microbiol 2004;70, DOI: 10.1128/AEM.70.2.943-953.2004.

3. Babu M, Beloglazova N, Flick R et al. A dual function of the CRISPR-Cas system in bacterial antivirus immunity and DNA repair. Mol Microbiol 2011;79, DOI: 10.1111/j.1365-2958.2010.07465.x.

4. Berkemer SJ, Maier LK, Amman F et al. Identification of RNA 3′ ends and termination sites in Haloferax volcanii. RNA Biol 2020;17, DOI: 10.1080/15476286.2020.1723328.

5. Bitan-Banin G, Ortenberg R, Mevarech M. Development of a gene knockout system for the halophilic archaeon Haloferax volcanii by use of the pyrE gene. J Bacteriol 2003;185, DOI: 10.1128/JB.185.3.772-778.2003.

6. Bligh E, Dyer W. A rapid method of total lipid extraction and pruification. Can J Biochem Physiol 1959;37, DOI: 10.1139/o59-099.PMID:13671378.

7. Boya BR, Kumar P, Lee JH et al. Diversity of the tryptophanase gene and its evolutionary implications in living organisms. Microorganisms 2021;9, DOI: 10.3390/microorganisms9102156.

8. Brendel J, Stoll B, Lange SJ et al. A complex of cas proteins 5, 6, and 7 is required for the biogenesis and stability of clustered regularly interspaced short palindromic repeats (CRISPR)-derived RNAs (crRNAs) in Haloferax volcanii. Journal of Biological Chemistry 2014;289, DOI: 10.1074/jbc.M113.508184.

9. Delpech F, Collien Y, Mahou P et al. Snapshots of archaeal DNA replication and repair in living cells using super-resolution imaging. Nucleic Acids Res 2018;46, DOI: 10.1093/nar/gky829.

10. Duggin IG, Aylett CHS, Walsh JC et al. CetZ tubulin-like proteins control archaeal cell shape. Nature 2015;519, DOI: 10.1038/nature13983.

11. Edelstein AD, Tsuchida MA, Amodaj N et al. Advanced methods of microscope control using μManager software. J Biol Methods 2014;1, DOI: 10.14440/jbm.2014.36.

12. Fahnert B, Lilie H, Neubauer P. Inclusion bodies: formation and utilisation. Adv Biochem Eng Biotechnol 2004;89, DOI: 10.1007/b93995.

13. Faure G, Makarova KS, Koonin E V. CRISPR–Cas: Complex Functional Networks and Multiple Roles beyond Adaptive Immunity. J Mol Biol 2019;431, DOI: 10.1016/j.jmb.2018.08.030.

14. Gaimster H, Cama J, Hernández-Ainsa S et al. The indole pulse: A new perspective on indole signalling in Escherichia coli. PLoS One 2014;9, DOI: 10.1371/journal.pone.0093168.

15. Ithurbide S, de Silva R, Brown H et al. A vector system for single and tandem expression of cloned genes and multi-colour fluorescent tagging in Haloferax volcanii. Microbiology (United Kingdom) 2024;170, DOI: 10.1099/mic.0.001461.

16. Jevtić Ž, Stoll B, Pfeiffer F et al. The Response of Haloferax volcanii to Salt and Temperature Stress: A Proteome Study by Label-Free Mass Spectrometry. Proteomics 2019;19, DOI: 10.1002/pmic.201800491.

17. Joffré E, Xiao X, Correia MSP et al. Analysis of Growth Phases of Enterotoxigenic Escherichia coli Reveals a Distinct Transition Phase before Entry into Early Stationary Phase with Shifts in Tryptophan, Fucose, and Putrescine Metabolism and Degradation of Neurotransmitter Precursors. Microbiol Spectr 2022;10, DOI: 10.1128/spectrum.01755-21.

18. Jones DL, Leroy P, Unoson C et al. Kinetics of dCas9 target search in Escherichia coli. Science 2017;357, DOI: 10.1126/science.aah7084.

19. Justice I, Kiesel P, Safronova N et al. A tuneable minimal cell membrane reveals that two lipid species suffice for life. Nat Commun 2024;15, DOI: 10.1038/s41467-024-53975-y.

20. Katewa SD, Katyare SS. A simplified method for inorganic phosphate determination and its application for phosphate analysis in enzyme assays. Anal Biochem 2003;323, DOI: 10.1016/j.ab.2003.08.024.

21. Kellermann MY, Yoshinaga MY, Valentine RC et al. Important roles for membrane lipids in haloarchaeal bioenergetics. Biochim Biophys Acta Biomembr 2016;1858, DOI: 10.1016/j.bbamem.2016.08.010.

22. Kim M, Seo DH, Park YS et al. Isolation of Lactobacillus plantarum subsp. Plantarum Producing C30 Carotenoid 4,4’-Diaponeurosporene and the Assessment of Its Antioxidant Activity. J Microbiol Biotechnol 2019;29, DOI: 10.4014/jmb.1909.09007.

23. Knight SC, Xie L, Deng W et al. Dynamics of CRISPR-Cas9 genome interrogation in living cells. Science 2015;350, DOI: 0.1126/science.aac6572.

24. Kopp J, Bayer B, Slouka C et al. Fundamental insights in early-stage inclusion body formation. Microb Biotechnol 2023;16, DOI: 10.1111/1751-7915.14117.

25. Landgraf D, Okumus B, Chien P et al. Segregation of molecules at cell division reveals native protein localization. Nat Methods 2012;9, DOI: 10.1038/nmeth.1955.

26. Large A, Stamme C, Lange C et al. Characterization of a tightly controlled promoter of the halophilic archaeon Haloferax volcanii and its use in the analysis of the essential cct1 gene. Mol Microbiol 2007;66, DOI: 10.1111/j.1365-2958.2007.05980.x.

27. Lee JH, Lee J. Indole as an intercellular signal in microbial communities. FEMS Microbiol Rev 2010;34, DOI: 10.1111/j.1574-6976.2009.00204.x.

28. Lestini R, Duan Z, Allers T. The archaeal Xpf/Mus81/FANCM homolog Hef and the Holliday junction resolvase Hjc define alternative pathways that are essential for cell viability in Haloferax volcanii. DNA Repair (Amst) 2010;9, DOI: 10.1016/j.dnarep.2010.06.012.

29. Liao Y, Ithurbide S, Evenhuis C et al. Cell division in the archaeon Haloferax volcanii relies on two FtsZ proteins with distinct functions in division ring assembly and constriction. Nat Microbiol 2021;6, DOI: 10.1038/s41564-021-00894-z.

30. Makarova KS, Aravind L, Grishin N V et al. A DNA repair system specific for thermophilic Archaea and bacteria predicted by genomic context analysis. Nucleic Acids Res 2002;30, DOI: 10.1093/nar/30.2.482.

31. Martens KJA, van Beljouw SPB, van der Els S et al. Visualisation of dCas9 target search in vivo using an open-microscopy framework. Nat Commun 2019;10, DOI: 10.1038/s41467-019-11514-0.

32. Martens KJA, Turkowyd B, Hohlbein J et al. Temporal analysis of relative distances (TARDIS) is a robust, parameter-free alternative to single-particle tracking. Nat Methods 2024, DOI: 10.1038/s41592-023-02149-7.

33. Di Martino P, Fursy R, Bret L et al. Indole can act as an extracellular signal to regulate biofilm formation of Escherichia coli and other indole-producing bacteria. Can J Microbiol 2003;49, DOI: 10.1139/w03-056.

34. Megaw J, Gilmore BF. Archaeal persisters: Persister cell formation as a stress response in Haloferax volcanii. Front Microbiol 2017;8, DOI: 10.3389/fmicb.2017.01589.

35. Movasaghi Z, Rehman S, Rehman IU. Fourier transform infrared (FTIR) spectroscopy of biological tissues. Appl Spectrosc Rev 2008;43, DOI: 10.1080/05704920701829043.

36. Noby N, Khattab SN, Soliman NA. Sustainable production of bacterioruberin carotenoid and its derivatives from Arthrobacter agilis NP20 on whey-based medium: optimization and product characterization. Bioresour Bioprocess 2023;10, DOI: 10.1186/s40643-023-00662-3.

37. Nuñez JK, Kranzusch PJ, Noeske J et al. Cas1-Cas2 complex formation mediates spacer acquisition during CRISPR-Cas adaptive immunity. Nat Struct Mol Biol 2014;21, DOI: 10.1038/nsmb.2820.

38. Nußbaum P, Gerstner M, Dingethal M et al. The archaeal protein SepF is essential for cell division in Haloferax volcanii. Nat Commun 2021;12, DOI: 10.1038/s41467-021-23686-9.

39. Nußbaum P, Kureisaite-Ciziene D, Bellini D et al. PRC domain-containing proteins modulate FtsZ-based archaeal cell division. Nat Microbiol 2024, DOI: 10.1101/2023.03.28.534543.

40. Ortenberg R, Rozenblatt-Rosen O, Mevarech M. The extremely halophilic archaeon Haloferax volcanii has two very different dihydrofolate reductases. Mol Microbiol 2000;35, DOI: 10.1046/j.1365-2958.2000.01815.x.

41. Ovesný M, Křížek P, Borkovec J et al. ThunderSTORM: A comprehensive ImageJ plug-in for PALM and STORM data analysis and super-resolution imaging. Bioinformatics 2014;30, DOI: 10.1093/bioinformatics/btu202.

42. Patro M, Duggin IG, Albers SV et al. Influence of plasmids, selection markers and auxotrophic mutations on Haloferax volcanii cell shape plasticity. Front Microbiol 2023;14, DOI: 10.3389/fmicb.2023.1270665.

43. Pende N, Sogues A, Megrian D et al. SepF is the FtsZ anchor in archaea, with features of an ancestral cell division system. Nat Commun 2021;12, DOI: 10.1038/s41467-021-23099-8.

44. Pérez-Arnaiz P, Dattani A, Smith V et al. Haloferax volcanii-A model archaeon for studying DNA replication and repair: Haloferax volcanii, a model archaeon. Open Biol 2020;10, DOI: 10.1098/rsob.200293.

45. Pinkard H, Stuurman N, Ivanov IE et al. Pycro-Manager: open-source software for customized and reproducible microscope control. Nat Methods 2021, DOI: 10.14440/jbm.2014.36.

46. Polívka T, Frank HA. Molecular factors controlling photosynthetic light harvesting by carotenoids. Acc Chem Res 2010;43, DOI: 10.1021/ar100030m.

47. Przykaza K, Woźniak K, Jurak M et al. Properties of the Langmuir and Langmuir–Blodgett monolayers of cholesterol-cyclosporine A on water and polymer support. Adsorption 2019;25, DOI: 10.1007/s10450-019-00117-2.

48. Rados T, Andre K, Cerletti M et al. A sweet new set of inducible and constitutive promoters for haloarchaea. Front Microbiol 2023, DOI: 10.3389/fmicb.2023.

49. Risović D, Frka S, Kozarac Z. Application of Brewster angle microscopy and fractal analysis in investigations of compressibility of Langmuir monolayers. Journal of Chemical Physics 2011;134, DOI: 10.1063/1.3522646.

50. Rizk S, Henke P, Santana-Molina C et al. Functional diversity of isoprenoid lipids in Methylobacterium extorquens PA1. Mol Microbiol 2021;116, DOI: 10.1111/mmi.14794.

51. Rolfsmeier ML, Laughery MF, Haseltine CA. Repair of DNA double-strand breaks following UV damage in three Sulfolobus solfataricus strains. J Bacteriol 2010;192, DOI: 10.1128/JB.00667-10.

52. Rollie C, Schneider S, Brinkmann A et al. Intrinsic sequence specificity of the Cas1 integrase directs new spacer acquisition. Elife 2015, DOI: 10.7554/eLife.08716.001.

53. Savery NJ. The molecular mechanism of transcription-coupled DNA repair. Trends Microbiol 2007;15, DOI: 10.1016/j.tim.2007.05.005.

54. Schindelin J, Arganda-Carreras I, Frise E et al. Fiji: An open-source platform for biological-image analysis. Nat Methods 2012;9, DOI: 10.1038/nmeth.2019.

55. Schmidt U, Weigert M, Broaddus C et al. Cell Detection with Star-convex Polygons. International Conference on Medical Image Computing and Computer-Assisted Intervention (MICCAI) 2018;11071, DOI: 10.1007/978-3-030-00934-2_30.

56. Seel W, Baust D, Sons D et al. Carotenoids are used as regulators for membrane fluidity by Staphylococcus xylosus. Sci Rep 2020;10, DOI: 10.1038/s41598-019-57006-5.

57. De Silva RT, Abdul-Halim MF, Pittrich DA et al. Improved growth and morphological plasticity of haloferax volcanii. Microbiology (United Kingdom) 2021;167, DOI: 10.1099/mic.0.001012.

58. Stracy M, Jaciuk M, Uphoff S et al. Single-molecule imaging of UvrA and UvrB recruitment to DNA lesions in living Escherichia coli. Nat Commun 2016;7, DOI: 10.1038/ncomms12568.

59. Turkowyd B, Balinovic A, Virant D et al. A General Mechanism of Photoconversion of Green-to-Red Fluorescent Proteins Based on Blue and Infrared Light Reduces Phototoxicity in Live-Cell Single-Molecule Imaging. Angewandte Chemie 2017;129, DOI: 10.1002/ange.201702870.

60. Turkowyd B, Müller-Esparza H, Climenti V et al. Live-cell single-particle tracking photoactivated localization microscopy of Cascade-mediated DNA surveillance. Methods Enzymol 2019;616, DOI: 10.1016/bs.mie.2018.11.001.

61. Turkowyd B, Schreiber S, Wörtz J et al. Establishing Live-Cell Single-Molecule Localization Microscopy Imaging and Single-Particle Tracking in the Archaeon Haloferax volcanii. Front Microbiol 2020;11, DOI: 10.3389/fmicb.2020.583010.

62. Vink JNA, Martens KJA, Vlot M et al. Direct Visualization of Native CRISPR Target Search in Live Bacteria Reveals Cascade DNA Surveillance Mechanism. Mol Cell 2020;77, DOI: 10.1016/j.molcel.2019.10.021.

63. Walsh JC, Angstmann CN, Bisson-Filho AW et al. Division plane placement in pleomorphic archaea is dynamically coupled to cell shape. Mol Microbiol 2019;112, DOI: 10.1111/mmi.14316.

64. Wang D, Ding X, Rather P. Indole can act as an extracellular signal in Escherichia coli. J Bacteriol 2001;183:4210–6.

65. Wang J, Li J, Zhao H et al. Structural and Mechanistic Basis of PAM-Dependent Spacer Acquisition in CRISPR-Cas Systems. Cell 2015;163:853.

66. Weigert M, Schmidt U, Haase R et al. Star-convex Polyhedra for 3D Object Detection and Segmentation in Microscopy. 2020 IEEE Winter Conference on Applications of Computer Vision (WACV) 2020, DOI: 10.1109/WACV45572.2020.9093435.

67. van Wolferen M, Pulschen AA, Baum B et al. The cell biology of archaea. Nat Microbiol 2022;7, DOI: 10.1038/s41564-022-01215-8.

68. Wörtz J, Smith V, Fallmann J et al. Cas1 and Fen1 Display Equivalent Functions During Archaeal DNA Repair. Front Microbiol 2022;13, DOI: 10.3389/fmicb.2022.822304.

