## Supplemental Data 1 for "A novel expression system for imaging single-molecule fluorescence in Haloferax volcanii WR806 enables visualization of Cas1 involvement in UV-light associated DNA repair"

**Supplementary Table S1 – S5**

**Supplementary Figure S1 – S5**

**Supplementary Table S1.** Strains used in this work.

| **Strain** | **Genotype** | **Needed Additives** | **Reference** |
|---|---|---|---|
| NEB® 5-alpha Competent *E. coli* (High Efficiency) | *fhuA2Δ(argF-lacZ)U169 phoA glnV44 Φ80Δ(lacZ)M15 gyrA96 recA1 relA1 endA1 thi-1 hsdR17* | - | New England Biolabs (Frankfurt am Main, Germany) |
| *dam⁻/dcm⁻* Competent *E. coli* | *ara-14 leuB6 fhuA31 lacY1 tsx78 glnV44 galK2 galT22 mcrA dcm-6 hisG4 rfbD1 R(zgb210::Tn10)* $Tet^S$ *endA1 rspL136* ($Str^R$) *dam13::Tn9* ($Cam^R$) *xylA-5 mtl-1 thi-1 mcrB1 hsdR2* | - | New England Biolabs (Frankfurt am Main, Germany) |
| *H. volcanii* WR806 | DS70-wildtype (ΔpHV2), Δ*pyrE2*, Δ*trpA*, Δ*leuB*, Δ*hdrB*, Δ*crtI* | Thymidine, uracil, tryptophan | (Turkowyd et al., 2020) |
| *H. volcanii* H119 | DS70-wildtype (ΔpHV2), Δ*pyrE2,* Δ*trpA*, Δ*leuB* | Tryptophan, uracil | (Allers et al., 2004) |

**Supplementary Table S2.** Oligonucleotides used in this work.

| **Oligonucleotide** | **Sequence 5' → 3'** | **Application** |
|---|---|---|
| screening_pUE001_for | AGTGAGCGAGGAAGCGGAAG | Verification of positive transformants |
| screening_pUE001_rev | GTGGCGAGAAAGGAAGGGAAG | Verification of positive transformants |
| sequencing_Dendra_rev | TCGCCTTCGATGACGAACGC | Verification of positive transformants |
| pTA231_cas1_dendra_fwd | ccatcgcatttttcggcgcgAAGCTTGGTACCGATATCGAATTCGATATCAAGC | Backbone amplification pUE001-cas1:Dendra2Hfx, pUE001 ftsZ1:Dendra2Hfx |
| pTA231_assembly_dendra_rev | GGATCCGAGCTCGCGGCC | Backbone amplification pUE001-cas1:Dendra2Hfx, pUE001 ftsZ1:Dendra2Hfx |
| cas1_Dendra_fwd | gcggccgcgagctcggatccGACTTCGACGACTACTTCGAC | Insert amplification cas1, ftsZ1 |
| cas1_Dendra_rev | CGCGCCGAAAAATGCGATG | Insert amplification cas1, ftsZ1 |
| pUE001-Dendra2Hfx_fwd | gcggacctattgcgcatatgaacacgccgggcatcaac | Vector amplification pUE001-Dendra2Hfx |
| pUE001-Dendra2Hfx_rev | CATATGCGCAATAGGTCCGC | Vector amplification pUE001-Dendra2Hfx |

**Supplementary Table S3.** Vectors used in this work.

| **Vector name** | **Features** | **Reference** |
|---|---|---|
| pTA231-p.Syn-Dendra2Hfx | p.syn promoter, *trpA*, *ampR,* encodes cytosolically expressed Dendra2Hfx, shuttle vector | (Turkowyd et al., 2020)<br>Addgene: #164660 |
| pTA962-Dendra2Hfx | p.tna promoter, *hdrB*, *pyrE*2, *ampR*, encodes cytosolically expressed Dendra2Hfx, shuttle vector | This work |
| pTA962-FtsZ1:Dendra2Hfx | p.tna promoter, *hdrB*, *pyrE*2, *ampR*, encodes FtsZ1:Dendra2Hfx fusion protein, shuttle vector | (Turkowyd et al., 2020) |
| pTA962-Cas1:Dendra2Hfx | p.tna promoter, *hdrB*, *pyrE*2, *ampR*, encodes Cas1:Dendra2Hfx fusion protein, shuttle vector | (Wörtz, 2022) |
| pUE001-Dendra2Hfx | p.tna promoter, *trpA*, *ampR,* encodes cytosolically expressed Dendra2Hfx, shuttle vector | This work<br>Addgene # 234669 |
| pUE001-Cas1:Dendra2Hfx | p.tna promoter, *trpA*, *ampR,* encodes Cas1:Dendra2Hfx fusion protein, shuttle vector | This work<br>Addgene # 234670 |
| pUE001-FtsZ1:Dendra2Hfx | p.tna promoter, *trpA*, *ampR,* encodes FtsZ1:Dendra2Hfx fusion protein, shuttle vector | This work<br>Addgene # 234671 |

**Supplementary Table S4** Parameters used in ThunderSTORM 1.3 for localization of single emitters

| | |
|---|---|
| Analysis filter | WaveletFilter (B-spline)<br>scale: 2.0<br>oder: 3 |
| Analysis detector | Local maximum<br>connectivity: 8<br>threshold: 1.4*std.(Wave.F1) |
| Analysis estimator | PSF: integrated gaussian<br>fitting redius: 3<br>method: max. likelihood<br>initial sigma: 1.6<br>full image fitting: false<br>mfaEnabled: false<br>nMax: 0<br>pVAlue: 0.0<br>keepSameIntensity: false<br>intensityInRange: false |
| Post processing | [] |
| Is 3D | false |
| Is set 3D | true |

**Supplementary Table S5** Parameters used in *swift v0.4.3* to determine trajectories of single molecules.

| | |
|---|---|
| diffraction_limit | 150 |
| directed_motion | false |
| exp_displacement | 400 |
| exp_noise_rate | 10 |
| max_blinking_duration | 2 |
| max_displacement | 2.5 x exp_displacement |
| max_displacement_pp | 3.5 x exp_displacement |
| max_log_complexity | 13 |
| max_memory | 500 |
| max_particle_count | 2 |
| max_subgraph_size | 50000 |
| p_bleach | 0.1 |
| p_blink | 0.001 |
| p_reappear | 0.5 |
| p_switch | 0.001 |
| precision | 40 |
| precision_z | 100 |
| pruning_base | 1.5 |
| pruning_rate | 0.2 |
| random_seed | 42 |
| threads | 12 |
| w_diffusion | 2 |
| w_dir_diffusion | 1 |
| w_immobile | 1 |

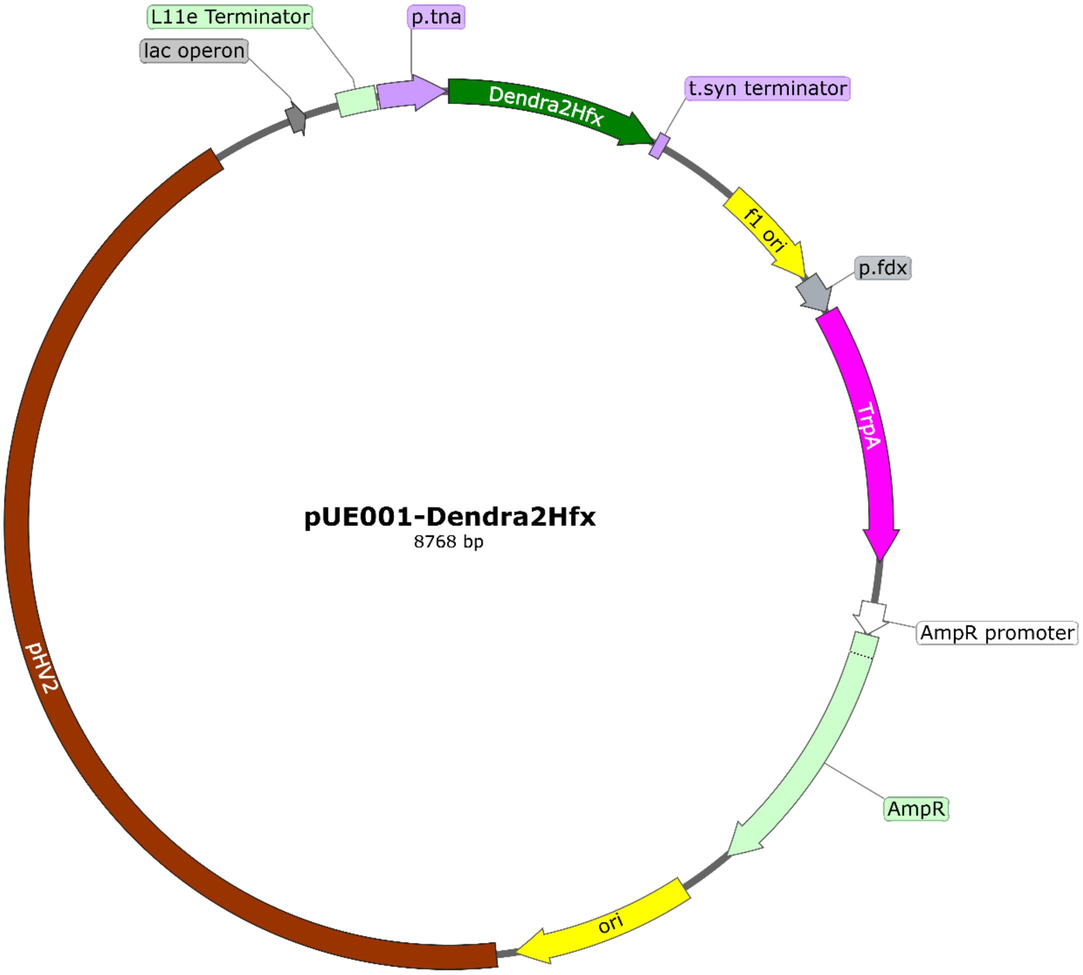


**Supplementary Figure S1.** Plasmid map of the novel expression system pUE001-Dendra2Hfx. pUE001 can be used as a shuttle vector in *E. coli* and *H. volcanii* and combines the selection marker trpA with the tryptophan inducible promoter p.tna.

Features for replication, selection and expression in *E. coli*: F1 ori: origin of replication; AmpR promoter: promoter for ampicillin resistance gene; *ampR*: ampicillin resistance gene for selection

Features for replication, selection, and expression in *H. volcanii: trpA*: tryptophan synthase encoding selection marker; p.fdx: constitutive promoter for the transcription of *trpA;* pHV2: origin of replication; p.tna: tryptophan inducible promoter for gene expression; L11e Terminator: rRNA terminator; t.syn: transcriptional terminator

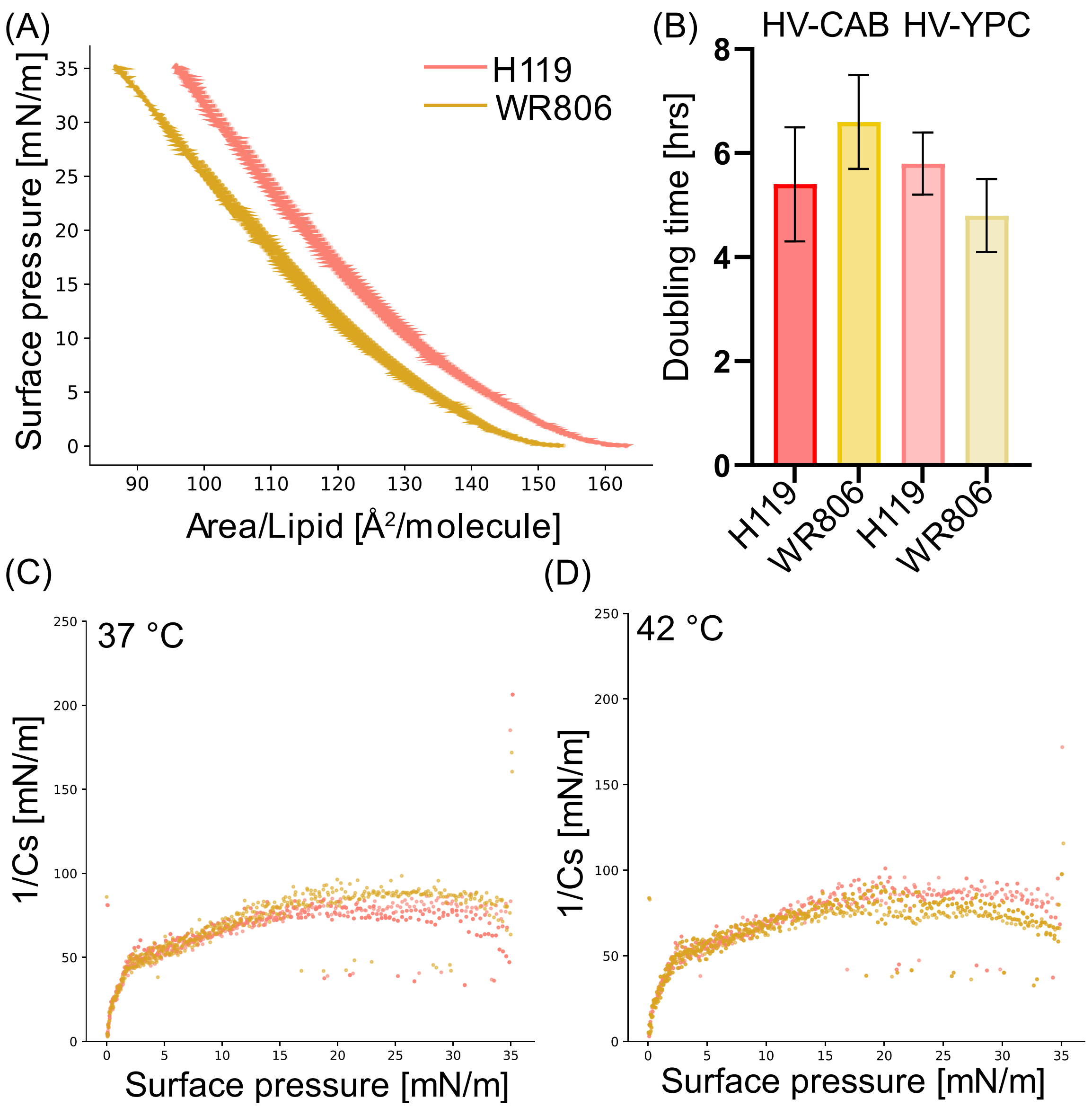


**Supplementary Figure S2.** Extended data for Figure 1 comparing strains H119 (red) and WR806 (yellow). (A) Area isotherms measuring surface pressure versus molecular area for lipid monolayers from extracted lipids from H119 and WR806 at 37 °C (biological duplicates, each with technical triplicates); (B) Doubling times of H119 and WR806 strains after transiting from HV-YPC to HV-CAB media or vice versa (means ± std. dev.; two biological replicates, each with three technical replicates). The indicated medium represents the final medium in which the strains were grown. (C, D) Compressibility modulus (k) of monolayer experiments performed with lipid extracts from H119 and WR806 at 37 °C (C) and 42 °C (D).

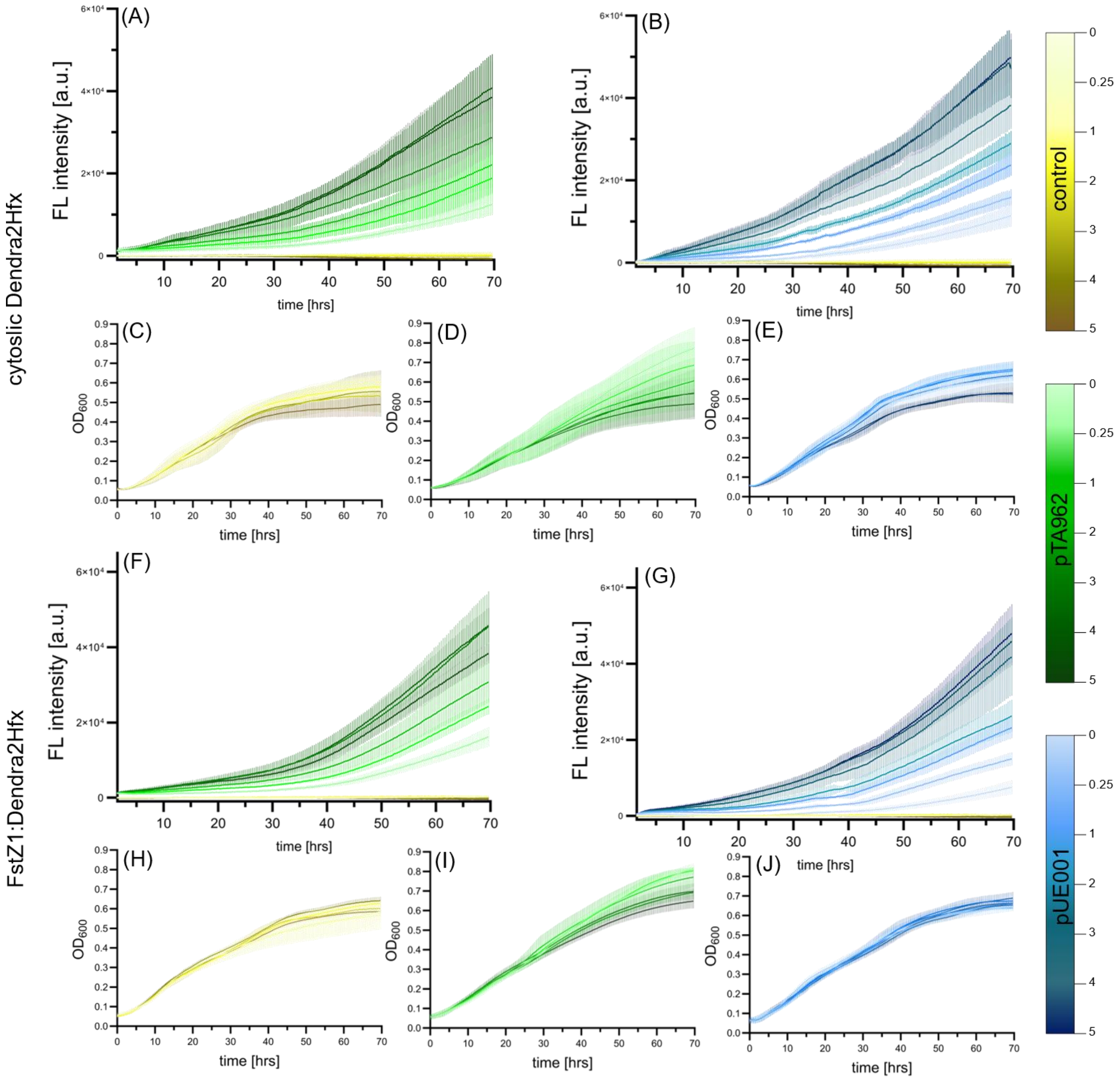


**Supplementary Figure S3**. Extended data to Figure 2. Fluorescence characterization of Dendra2Hfx expression in *H. volcanii* strain WR806. Raw fluorescence and OD measurements belonging to the normalized fluorescence intensities shown in Fig. 2A, B, D and E. Experiments were performed at 42°C in biological triplicates, each with technical duplicates, data is visualized as mean with standard deviation. Cultures carrying pTA962 (green) and pUE001 (blue) plasmids were induced with tryptophan concentrations of 0, 0.25, 1, 2, 3, 4, and 5 mM. Plasmid-free WR806 (yellow) served as negative control.

(A, B, F, G) Time-course fluorescence measurements of tryptophan-induced protein expression (ex 470 nm; em 501-519 nm, optimized for Dendra2Hfx green fluorescence). (A, B) Cytosolic Dendra2Hfx expression. (F, G) FtsZ1:Dendra2Hfx fusion protein expression.

(C, D, E, H, I, J) Measurement of optical density was determined by light absorption at a wavelength of 600 nm. (C, D, E) optical density measurements of WR806 (yellow), WR806 pTA962-Dendra2Hfx (green), and WR806 pUE001-Dendra2hfx (blue). (H, I, J) Optical density measurements of WR806 (yellow), WR806 pTA962-FtsZ1:Dendra2Hfx (green), and WR806 pUE001-FtsZ1:Dendra2hfx (blue).

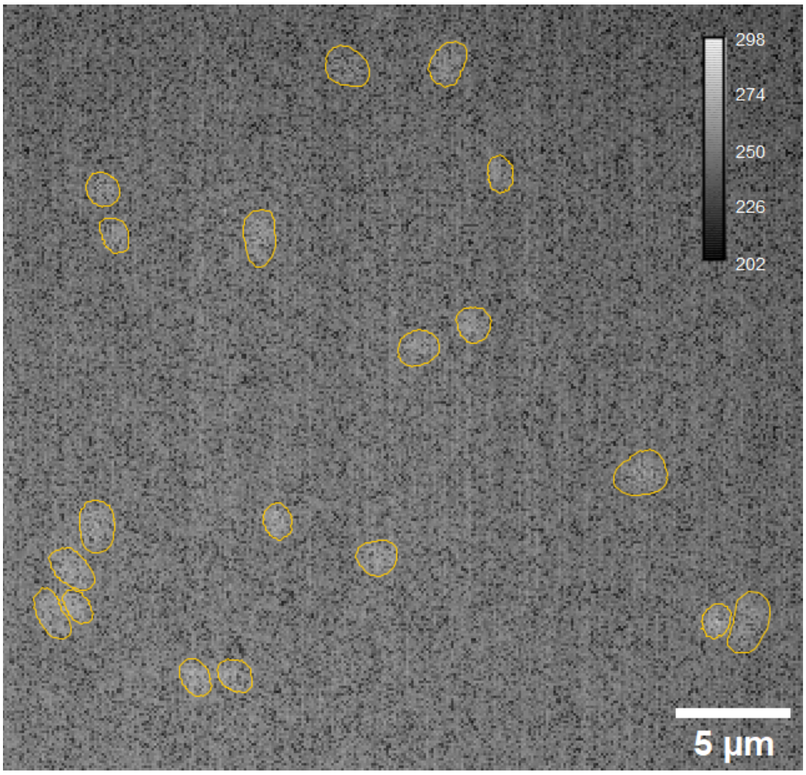


**Supplementary Figure S4.** The fluorescence of green Dendra2Hfx in WR806 pTA231-FtsZ1:Dendra2Hfx at OD 1 was measured. Although the strain carried the correct plasmid as verified by colony PCR, no visible expression of the fusion protein FtsZ1-Dendra2Hfx was detected. This finding is consistent with earlier, unpublished observations during tests of fusion constructs for plasmids described in (Turkowyd et al., 2020). While cytosolically expressed fluorescent proteins were clearly visible (Turkowyd et al., 2020), fusion proteins in a pTA231 background consistently failed to show expression, as observed in our unpublished work. We attribute this suppression to the potential strong regulation of constitutively expressed transcripts that include native *H. volcanii* gene sequences. Having carefully confirmed these old preliminary results with above findings, we are now confident in the validity of the negative results and have decided to publish them.

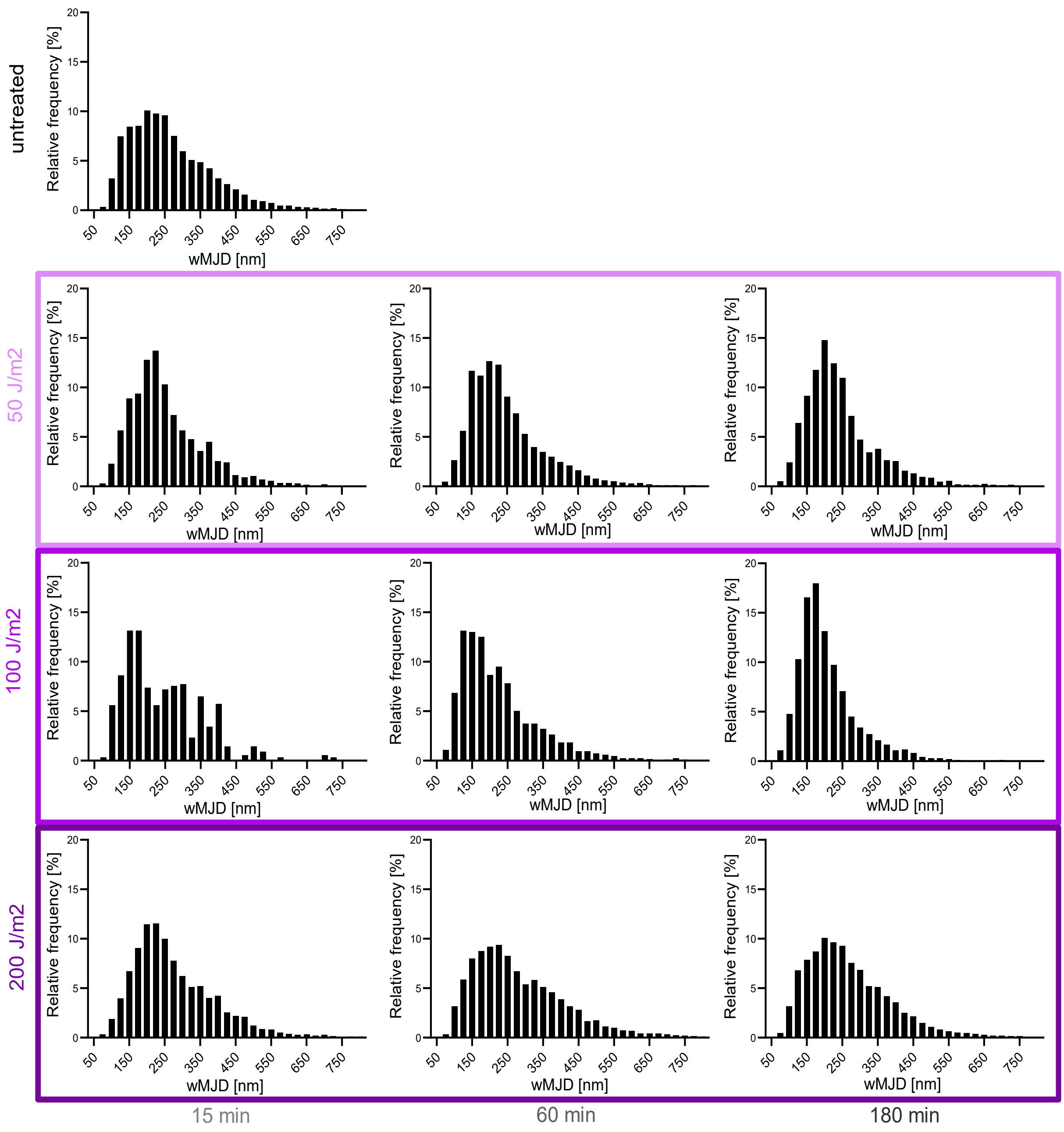


**Supplementary Figure S5.** Extended data to Figure 4. Analysis of Cas1-Dendra2Hfx single-molecule dynamics in response to UV-light damage in *H. volcanii* strain WR806 pUE001 as measured by single-particle tracking microscopy. Exponentially growing cells cultured without tryptophan were sub-cultured with 0.25 mM tryptophan for inducing Cas1 protein expression overnight. Cells were reinoculated under the same conditions to OD 0.1 and after 2 hours of growth, exposed to UV-light radiation at 265 nm at 0, 50, 100, or 200 J/m² and measured after a recovery time of 0, 15, 60, and 180 minutes post-exposure. Dynamics are quantified by weighted mean jump distances (wMJD) from individual single-molecule trajectories across two biological replicates. Mean jump distances were weighted by the respective number of localizations per trajectory. Light to dark purple boxes indicate the increased UV dosage and histograms are sorted according to the recovery time of the cells after exposure. The untreated control is shown in the upper row.

### References

Allers, T., Ngo, H. P., Mevarech, M., & Lloyd, R. G. (2004). Development of Additional Selectable Markers for the Halophilic Archaeon Haloferax volcanii Based on the *leuB* and *trpA* Genes. *Applied and Environmental Microbiology*, *70*(2), 943–953. https://doi.org/10.1128/AEM.70.2.943-953.2004

Turkowyd, B., Schreiber, S., Wörtz, J., Segal, E. S., Mevarech, M., Duggin, I. G., Marchfelder, A., & Endesfelder, U. (2020). Establishing Live-Cell Single-Molecule Localization Microscopy Imaging and Single-Particle Tracking in the Archaeon *Haloferax volcanii*. *Frontiers in Microbiology*, *11*. https://doi.org/10.3389/fmicb.2020.583010

Wörtz, J. E. (2022). *Haloferax volcanii:* Untersuchung alternativer Funktionen der Endonuklease Cas1 über die CRISPR-Cas Immunabwehr hinaus. Ulm University.
